# Optimized double emulsion flow cytometry with high-throughput single droplet isolation

**DOI:** 10.1101/803460

**Authors:** Kara K. Brower, Catherine Carswell-Crumpton, Sandy Klemm, Bianca Cruz, Gaeun Kim, Suzanne G.K. Calhoun, Lisa Nichols, Polly M. Fordyce

**Affiliations:** Department of Bioengineering, Stanford University, Stanford, California, USA; Chem-H Institute, Stanford University, Stanford, California, USA; Center for Molecular and Genetic Medicine, Stanford University, Stanford, California, USA; Department of Genetics, Stanford University, Stanford, California, USA; Department of Physics, California State Polytechnic Institute, Pomona, CA, USA; Department of Chemical Engineering, Stanford University, Stanford, California, USA; Chan-Zuckerberg Biohub, San Francisco, CA, USA

## Abstract

Droplet microfluidics has made large impacts in diverse areas such as enzyme evolution, chemical product screening, polymer engineering, and single-cell analysis. However, while droplet reactions have become increasingly sophisticated, phenotyping droplets by a fluorescent signal and sorting them to isolate variants-of-interest remains a field-wide bottleneck. Here, we present an optimized double emulsion workflow, sdDE-FACS, that enables high-throughput phenotyping, selection, and sorting of droplets using standard flow cytometers. Using a 130 *μ*m nozzle, we demonstrate robust post-sort recovery of intact droplets, with little to no shear-induced droplet breakage, at high sort frequency (12-14 kHz) across two industry-standard FACS instruments. We report the first quantitative plate statistics for double emulsion droplet isolation and demonstrate single droplet recovery with >70% efficiency. In addition, we establish complete downstream recovery of nucleic acids from single, sorted double emulsion droplets, an advance in droplet sorting comparable with the capabilities of single-cell FACS. This work resolves several hurdles in the field of high-throughput droplet analysis and paves the way for a variety of new droplet assays, including rare variant isolation and multiparameter single-cell analysis, marrying the full power of flow cytometry with droplet microfluidics.

## 1 Introduction

Microfluidic droplet generation is a powerful technique for encapsulating biological molecules or cells within precisely controlled nL- to pL- volumes, making it possible to perform up to 10^7^ reactions in parallel with low per-reaction costs ^1^. Microfluidic droplets have been used for a wide variety of applications, including directed evolution of enzymes and proteins ^2–6^, digital PCR ^7^, large-scale gene assembly ^8^, cell culture ^9,10^, and, recently, single-cell genomic, epigenomic, and transcriptomic analyses ^11–15^. In the past ten years, droplet technologies have been translated to a variety of commercial assays (*e.g.*, ddPCR, Bio-rad; Chromium, 10X Genomics), representing perhaps the largest commercial adoption of microfluidic technologies to-date. However, while the number of possible reactions within droplet microreactors has increased, screening, sorting, and isolating sub-populations of droplets for downstream processing remains technically challenging ^16,17^.

Fluorescent readouts in droplet assays allow for quantitative measurement of reaction progress and outcome ^18–21^. When combined with an ability to sort droplets by their fluorescence, droplets can be binned by one or more signals and their nucleic acid content analyzed to identify variants responsible for activity (the *genotype-to-phenotype* linkage) ^2,5,7,22^. Currently, fluorescence-activated droplet sorting (FADS) remains the most common approach for droplet analysis and sorting ^16^. FADS and other variants of the technique (*e.g.* flow dropometry (FD) and picodispersion) analyze and sort water-in-oil (W/O) droplets based on fluorescence using a microfluidic chip with embedded electrodes and an associated optical assembly (for dielectrophoretic sorting and droplet imaging, respectively) ^19,23–25^. While FADS allows accurate droplet screening, high accuracy sorting is limited to slow sorting rates (0.1-2 kHz), only 1 or 2 fluorescence channels can be probed simultaneously, and downstream sorting requires extensive additional equipment ^19^. In addition, FADS requires custom devices and instruments that are technically demanding to build, maintain, and operate, limiting adoption to a few laboratories worldwide ^19,21,24^. Finally, demonstration of single-droplet isolation and sorting via FADS has been limited; custom fabricated machinery is required and no automation capabilities are currently available ^26^.

Fluorescence-activated cell sorting (FACS) instruments provide an appealing alternative to FADS for droplet sorting. FACS cytometers boast excellent signal discrimination and sensitivity ^27–30^, unparalleled multi-parameter analysis capabilities (2-18 fluorescence channels) ^31–35^, and established workflows for robust plate-based deposition of cells ^36–41^. FACS instruments are integrated, easy-to-operate, and widely available at most institutions. As a result, FACS remains the most ubiquitous and pervasive technique for cellular phenotyping worldwide ^42,43^. FACS allows quantitative measurement of cell surface markers and intracellular proteins with subsequent gating to isolate and enrich for subpopulations of interest, including rare variants, for down-stream processing (*e.g.*, sequencing, qPCR, clonal expansion) ^30^. The landmark demonstration of FACS for the detection, sorting, and nucleic acid recovery from individual, single cells ushered in a new era of single-cell analysis ^37,44^, allowing high-throughput investigation of the linkage between genotype and phenotype in each cell for many cells in parallel.

The ability to sort single droplets would be equally transformative in extending the capabilities of current single-variant droplet encapsulation techniques. However, sorting via FACS requires the ability to charge the aqueous fluid surrounding particles so that targets of interest can sorted by electrostatic deflection ^45^. Standard water-in-oil (W/O) droplets typically used for FADS are incompatible with FACS, as the insulating oil surrounding the aqueous core of W/O droplets is immiscible with the aqueous sheath fluids used in flow cytometers ^46^. Translating FACS to use with droplets therefore requires more complex water-oil-water (W/O/W) double emulsion (DE) droplets.

DEs have a double-droplet architecture in which an inner aqueous core (similar to typical single emulsions used in FADS) are encapsulated in an outer oil shell that is subsequently surrounded by aqueous fluid ^20,47^. DE droplets can therefore be suspended in an aqueous buffer that can be mixed with standard FACS sheath buffers (*e.g.* PBS). Prior work has established that FACS instruments can detect and sort DE droplets ^2,5,7,7,20,46,48–50^, but post-sort droplet recovery has been poor (~40-70% droplet survival post-FACS with a large fraction of ruptured droplets in post-sort images ^7^). Droplet rupture likely results from shear-induced droplet breakage during FACS and can lead to significant microreactor cross-contamination ^7^. As a result, downstream nucleic acid recovery from double emulsions, especially at low droplet numbers, has been inefficient or unsuccessful ^2,5,48^. Moreover, no technique to date has been able to reliably isolate individual DEs. High-throughput, individual droplet isolation would unlock possibilities for single-cell and rare-variant assays, as well as subpopulation enrichment at low droplet numbers, allowing direct investigation of *genotype-to-phenotype* linkages without cross-contamination.

Here, we demonstrate an improved workflow for double emulsion FACS (sdDE-FACS) that allows high-throughput, quantitative phenotyping and sorting of DE droplets with high rates of recovery, down to single droplet isolation. Using optimized surfactant mixtures, we generate stable DE droplets capable of withstanding the shear forces needed for FACS and any required thermocycling without detectable breakage. In addition, we describe droplet preparation and FACS settings for reliable, clog-free sorting with robust post-sort recovery across multiple instruments. sdDE-FACS provides high dynamic range and signal discrimination, with demonstrated DE sorting efficiencies of ~ 60-70% efficiency (near-complete droplet survival) and over 97% target specificity, on par with the capabilities of single cell FACS ^36,43,51–53^. Finally, we demonstrate robust recovery of nucleic acids from DE droplets via qPCR, with no evidence of well-to-well cross-contamination. To our knowledge, this represents the first demonstration of high-efficiency DE droplet sorting and complete nucleic acid recovery via FACS to the level of single droplet isolation.

These new capabilities significantly expand the available repertoire of droplet reactions to allow high-throughput phenotypic screening, reliable rare variant enrichment, and reduced assay costs. sdDE-FACS paves the way for a wide variety of new *genotype-to-phenotype* droplet assays by combining the throughput of droplet microfluidics with the power of single-cell FACS.

## 2 Results and Discussion

### 2.1 DE-FACS Workflow and Pipeline

To enable high-throughput sorting and analysis of DE droplet populations via FACS, we developed and optimized a 3-stage pipeline (<u>s</u>ingle <u>d</u>roplet <u>D</u>ouble <u>E</u>mulsion <u>FACS</u>-sdDE-FACS, **Fig. 1**). During the first stage (DE droplet library generation, **Fig. 1A**), variant libraries (*e.g.* prokaryotic or eukaryotic cells, nucleic acids, or proteins) are encapsulated within the inner aqueous phase of DE droplets in accordance with a Poisson-distributed occupancy to ensure most droplets are either empty or contain a single variant. Each variant is co-encapsulated with any required assay reagents (*e.g.* enzymes, buffers, dyes, or antibodies) and surfactants to stabilize the W/O/W droplet architecture ^7,54,55^; thermocycling steps for RT, PCR, incubation or other reactions can be performed at this stage as needed.

**Fig. 1.**
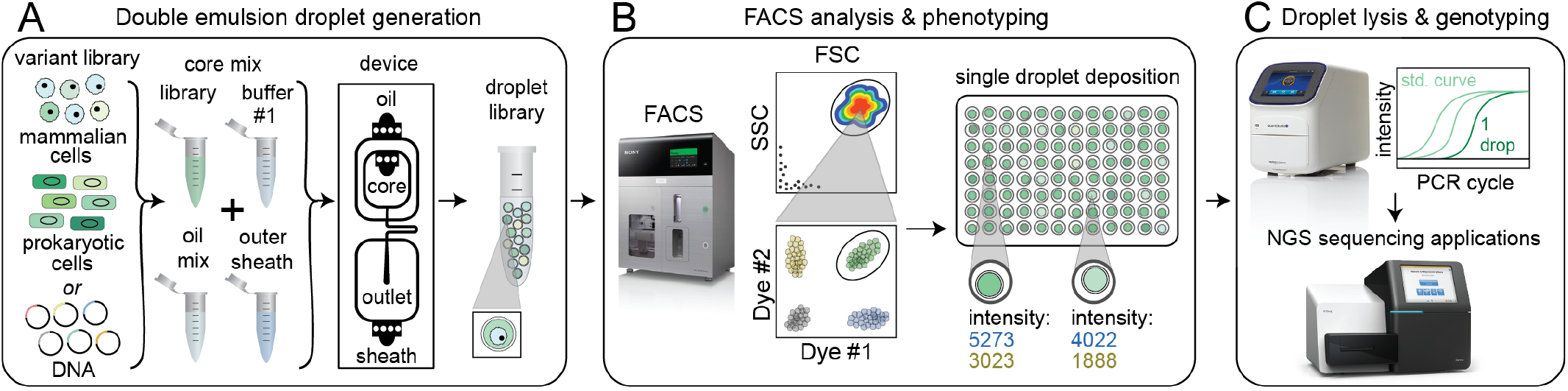
sdDE-FACS workflow. **(A)** Cell or DNA variants of interest are loaded into a DE droplet generator device to produce a library of droplets each containing a different variant. DEs can be generated for a wide variety of reactions by adjusting core mix reagents and buffers, number of core inlets, and droplet size. **(B)** DE droplets are analyzed via FACS to quantify morphology (FSC vs. SSC) and relevant fluorescence signals (by fluorescent intensity) and then sorted into wells of a multiwell plate. **(C)** Sorted DE droplets can be lysed to recover nucleic acids for downstream applications, such as qPCR or next-generation sequencing, to determine the droplet genotype responsible for a particular phenotype (*e.g.* enzymatic reaction turnover, presence of a specific cell type, or completion of a cellular reaction).

In the next stage (FACS phenotyping, **Fig. 1B**), DE droplets are quantitatively analyzed based on size and fluorescence and sorted via FACS into designated wells of a multi-well destination plate or other sort vessel. Sorting can either collect many DE droplets from a desired population or deposit *individual* droplets into particular wells, thereby directly linking DE droplet phenotype to an output plate well location. After sorting, pools or individual DE droplets can be lysed and processed downstream via various plate-based reaction techniques, including qPCR and next-generation sequencing (DE droplet genotyping, **Fig. 1C**). Barcodes linking DE droplets to well position can be incorporated at this stage to retain information linking nucleic acid sequences to their well location, thereby preserving *genotype-to-phenotype* linkages during subsequent pooled processing.

### 2.2 Double emulsion droplet generation device

Quantitative, high-throughput DE droplet phenotyping via FACS requires generation of highly monodisperse and stable DEs. For successful sorting, DE droplets must be significantly smaller (10-50 *μ*m in diameter) than commercial FACS nozzles (typically 70-130 *μ*m in diameter) while simultaneously large enough to encapsulate variants of interest (0.005-3 pL for bacteria to large mammalian cells, respectively) within the inner core volume (2-50 pL) ^7,46^. Smaller droplet sizes lower droplet deformation during FACS and thus minimize likelihood of DE droplet breakage ^55^.

To generate FACS-compatible DE droplets, we fabricated a one-step microfluidic dual-flow focusing device for W/O/W droplet generation based on previously-published designs ^56,57^ **(Fig. 2A, Fig. S1)**. This device is easy and inexpensive to operate, requiring only syringe pumps and a low-cost microscope with a high-speed camera to visualize droplets within the device (**Fig. S1, Table S1**). Devices were designed to produce W/O/W droplets significantly smaller than typical FACS nozzles (~30*μ*m, <5% CV), with channel heights of 15 *μ*m for the inner aqueous and oil phases (first flow focuser for W/O droplet generation) and 40 *μ*m for outer aqueous phase (second flow focuser to wrap the oil shell with aqueous buffer and create the W/O/W droplet). Larger or smaller double emulsions can be generated with scaled versions of this device; we have generated DE droplet populations from 27.63 - 48.36 *μ*m (see *Supplemental Information*), all of which perform well with this workflow. However, any custom or commercial device can be used to generate DE droplets compatible with sdDE-FACS as long as polydispersity is minimized (droplet CV <20%). Large size variation of the droplet sample, which is normally concomitant with the presence of significant free oil, increases the chance of clogging during FACS.

**Fig. 2.**
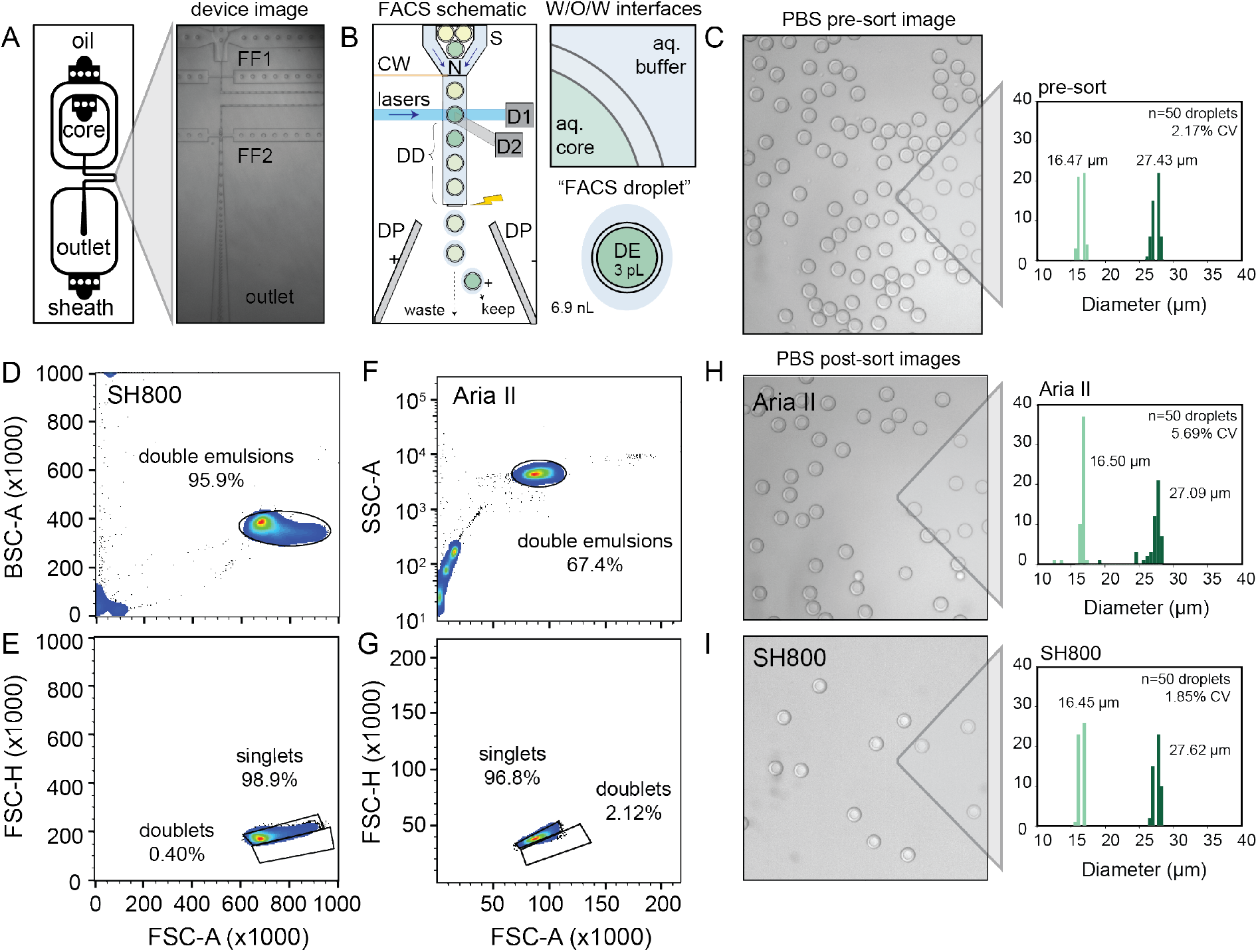
DE droplet analysis via sdDE-FACs. **A** Schematic of dual flow-focuser (FF1, FF2) DE droplet generator and image of DE droplet generation. **B** Schematic of DE droplet manipulations during FACS. DE droplets pass through a nozzle (N) and are hydrodynamically focused by sheath flow (S) prior to interrogation via lasers within the flow cell. Signals are read out via detectors (examples: D1, D2). After a specified droplet delay (DD), the sheath stream is charged via a charge wire (CW) to charge an individual FACS droplet containing a DE-of-interest prior to break-off; charged droplets are then deflected to a specific well (keep) or waste (W) via fields generated between dielectric plates (DP). Insets show a “FACS droplet” with encapsulated DE (respective volumes of each droplet indicated) as well as associated surfactant-stabilized W/O/W interfaces to recover DEs-of-interest. **C** Representative image of pre-sort DE droplets and pre-sort size distributions (light green = inner diameter, dark green = outer diameter, CV = variation of total diameter). **D,F** FACS light scatter gates of DEs on the SH800 and Aria II, respectively. (25,000 total events visualized, randomly sampled). A 9.9K threshold was applied to SH800 data eliminate small particulates/electronic noise/debris from gating. **E,G** Daughter singlet gates of the parental double emulsion populations per sorter. **H** Image and size distribution for DE droplets after sorting with the Aria II. **I** Image and size distribution for DE droplets after sorting with the SH800. (n=50 droplets analyzed for all size histograms).

### 2.3 Double emulsion surfactant selection

DE stability is critical to robust performance of sdDE-FACS. Appropriate surfactant choice in the aqueous and oil phases is required to stabilize the inner and outer water-oil boundaries of DEs throughout droplet formation, storage, and reaction processing ^46,47,54^. FACS sorting of DE droplets poses an even greater challenge, as DE droplets must remain intact even when exposed to high flow rates and shear forces (**Fig. 2B**) ^55^. During FACS, DE droplets are diluted in a diluent suspension buffer and loaded into the instrument, where they meet a fast-moving sheath buffer (with typical flow pressures of 1-10 psi). This sheath flow hydrodynamically focuses DE droplets and carries them into a flow cell where they are excited by a series of lasers for quantitative phenotyping ^27,45^. After laser interrogation, the sheath flow is acoustically vibrated to create a stable breakoff of water-in-air droplets (“FACS droplets”, 6.9 nL for a 130 *μ*m nozzle) that encapsulate DE droplets for sorting (now a triple-droplet architecture). Immediately prior to break-off, the sheath fluid is charged (by the charge wire, CW, **Fig. 2B**) if a droplet-of-interest meeting a phenotypic gate criteria is detected. This charge is imparted on the aersolized DE-containing “FACS droplet” for electrostatic deflection into wells. Only targeted droplets receive a charge; DEs not meeting the selected gate are directed to waste. The charge timing is decided by an important sorting parameter, called droplet delay (DD, **Fig. 2B**). Droplet delay is the timing between laser interrogation of an event of interest and its presence at the drop break-off point, which is impacted by particle size and hydrodynamics in the sort stream ^45^. Droplet delay must be determined empirically for reliable sorting. Robust recovery of DE droplets requires that the vast majority of droplets remain intact throughout this process to prevent cross-contamination (*e.g.* through breakage in the flow stream or FACS droplet spray).

Surfactants stabilize DE droplets by decreasing interfacial tension and distributing charge density ^54,55^. Surfactants locally crowd, absorb or “skin” at droplet oil-water and water-oil interfaces within the DE itself, between the DE and flow stream, and between the DE and larger “FACS droplet” to stabilize the droplet prior to and during FACS (**Fig. 2B**) ^7^. Prior work sorting double or single emulsions via FACS and FADS, respectively, have employed a wide variety of surfactants **(Table S2)**. Based on these reports, we selected a fluorinated oil (HFE 7500) and ionic surfactant (PEG-Krytox FS-H 157) combination previously shown to exhibit excellent biocompatibility, no leakage between phases, and high stability under storage and reaction thermocycling ^7,10,54^. After empirical testing, we modified this recipe as follows: (1) to reduce DE core droplet deformation under shear, we decreased inner and outer aqueous phase viscosities (which lowers viscous stress) and reduced inner aqueous phase non-ionic surfactant concentration, as well as lowered FACS sheath flow rates and increased nozzle size, as recommended by prior experimental ^7^ and theoretical work ^55^; and (2) to reduce FACS stream instability and clogging, we increased carrier aqueous phase non-ionic surfactant concentrations (which lowers shear and appears to prevent satellite oil formation during droplet generation), and reduced overall surfactant in the FACS diluent buffer ^55^. Finally, we osmotically balanced the inner and outer aqueous phases during droplet manipulation to prevent osmotic droplet lysis ^7^. This final formulation (**Table 1**) has yielded highly monodisperse (CV <5%) DE droplets across hundreds of samples. For cellular applications, 0.1-1% Tween-20 can be replaced with 0.5-2% BSA without loss of DE stability. A typical population (**Fig. 2C**) has mean diameters of 16.47 ± 0.47 *μ*m and 27.43 ± 0.60 *μ*m for the inner core and total droplet diameter, respectively (mean ± standard deviation) with an overall CV of 2.17% **(Fig. 2C)**. DE droplets can be stored for months to years without significant size changes, are compatible with a wide range of internal reagents, and are stable under reaction incubation and thermocycling.

**Table 1.**
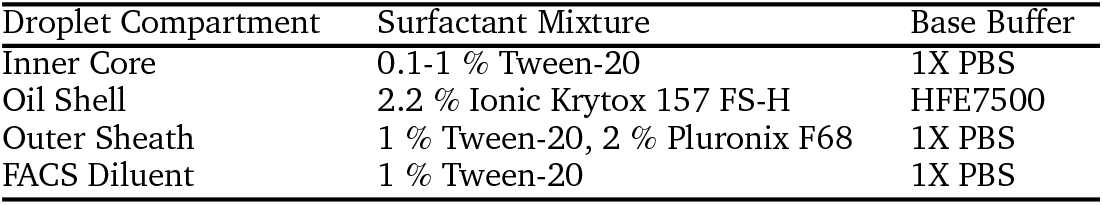
Double emulsion surfactant mix for stable droplet generation and FACS recovery. Base buffers can be substituted as desired given aqueous phases are osmotically-matched.

### 2.4 Droplet phenotyping using DE-FACS

Next, we tested the ability to visualize and sort DE droplet populations on two widely available FACS instruments, an Aria II (Becton Dickinson, BD) and a SH800 (Sony Biotechnologies) **(Table 2)**. The Aria II couples piezoelectric droplet generation with a traditional quartz cuvette for sample interrogation using high-powered lasers for excitation and collection optics gel-coupled to the cuvette for emission; it represents the current literature standard for droplet flow cytometry. The SH800 instead uses a microfluidic approach, where sample fluid channels, the laser interrogation window, and the sorting nozzle are integrated on a disposable chip; sample excitation uses lower-powered lasers and emission optics utilize just a single optical fiber without gel-coupling. In contrast with the Aria II, the Sony SH800 is relatively low cost, easy to operate, and requires minimal training.

**Table 2.** Comparison of FACS sorter instruments

| FACS Parameter | Aria II (BD) | SH800 (Sony) |
| --- | --- | --- |
| Laser Interrogation Vessel | Rectangular Cuvette | Microfluidic Chip |
| Emission Optic Coupling | Gel-coupled | Air |
| Laser Excitation | Spatially Separate | Colinear |
| Laser Power | 50-200 mW | 30 mW |
| Acoustic Mechanism | Up-down | Side-side |
| Nozzle | Ring | Chip-Integrated |
| ND Attenuation | Yes | No |
| Ease-of-Use | Requires expert training | Simple |

DE droplets are larger and more deformable than typical cells ^55^, requiring significant optimization of FACS instrument settings (*e.g.* scatter thresholds, laser gains, flow pressures, droplet harmonics, and droplet delays) to ensure consistent and quantitative detection and sorting **(Table 3)**. While previous DE droplet sorting used small (70 *μ*m or 100 *μ*m) sort nozzles ^7,20,46^, we employed a large (130 *μ*m) sort nozzle for sdDE-FACS to minimize droplet shear and breakage. The 130 *μ*m nozzle is a standard nozzle size in both instruments typically used for large cell types in FACS.

**Table 3.**
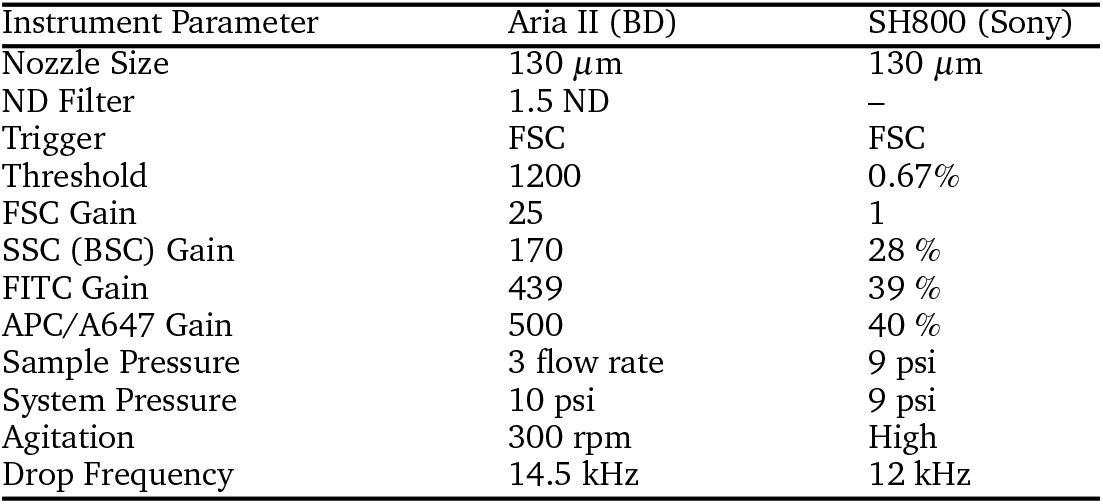
Optimized FACS Instrument Parameters for 30-50 *μ*m double emulsion droplet analysis and high-recovery sorting

With these optimized parameters, forward versus side scatter (FSC *vs.* SSC) distributions on the Aria II and the SH800 for the same population of DEs revealed distinct, tight clusters of DE droplets with limited scatter within 2 orders of magnitude (**Fig. 2D,F**). Compared to the previous Aria II literature benchmark of 43.9% for FSC *vs.* SSC purity (10,000 events, with similar gating strategy ^46^), this represents an overall increase in throughput of >25% on the Aria II **(Fig. 2D,F)**. More importantly, a extraneous scatter on both instruments is significantly decreased, indicating less droplet breakage and free oil with sdDE-FACS ^7,20,46^. Bivariate plots of double emulsion forward scatter height versus area (FSC-H *vs.* FSC-A, **Fig. 2E,G**) provide discrimination of single droplets from doublets or larger clusters; presence of a single dominant event cluster demonstrates that nearly all gated events are comprised of single DE droplets (doublet rates <3% on both instruments), suggesting that uniform, non-aggregate DEs are processed via sdDE-FACS, as verified by microscopy **(Fig. 2E,G,H,I)**. These results replicate across samples **(Fig. S2)**.

Images of DE droplets post-sort establish that populations remain monodisperse (Aria II: inner and total diameters of 16.50 ± 0.80 *μ*m and 27.08 ± 0.51 *μ*m, respectively; SH800: inner and total diameters of 16.45 ± 0.35 *μ*m and 27.61 ± 0.52 *μ*m, respectively (mean ± standard deviation)) with little breakage **(Fig. 2H,I)**. Post-sort DE size CV was lower for the SH800 (1.85% CV) as compared to the Aria II (5.69%), likely due to lower shear forces inside the microfluidic chip and integrated sort nozzle used on the SH800. Compared to the pre-sort population (2.17% CV, 27.43 *μ*m total diameter), the Aria II (5.69% CV, 27.08 *μ*m total diameter) post-sort populations yielded increased droplet oil shell size reductions. Further, accompanying oil droplets, possibly indicating droplet breakage, were more frequently observed with the Aria II post-sort than the SH800 **(Fig. 2H,I)**, likely due to adverse effects of higher shear in the Aria II. However, effects were relatively minor and did not significantly impact single droplet isolation in optical plate sorting or nucleic acid recovery. Interesting, the SH800 observed a droplet event lag time **(Fig. S3)**, perhaps due to packing of highly deformable DEs within flow chip before droplets were metered under laminar flow.

These results are representative across all 20+ droplet populations analyzed in this paper, extensible to different size DEs (**Figs. S4,S5,S6**) and are consistent for droplets having undergone additional pre-processing such as incubation and thermocycling. Overall performance was similar between the 2 instruments, and neither sorter clogged or paused during sorting. However, the SH800’s modularity, ease-of-use, gate purity, and excellent post-sort recovery are particularly well-suited to high-throughput applications.

### 2.5 Assessment of FACS dynamic range and limit of detection of double emulsions

FACS-based cell screening applications typically detect fluorescence emitted by cells containing fluorescent reporters with a dynamic range of ~4 orders of magnitude ^32,43,58^. To quantify the dynamic range and lower limit of detection for sdDE-FACS, we loaded DE droplets with five concentrations (0.01, 0.1, 1, 10, and 100 *μ*g/mL) of either FITC-labeled or Alexa-Fluor 647-labeled bovine serum albumin (BSA) and quantified emitted fluorescence via FACS on both instruments (**Fig. 3B,C,E,F**). Brightfield images confirmed that DE droplets remained highly monodisperse when loaded with dye-labeled BSA **(Fig. 3A,D)**. Measured intensities for DE droplets (gated by FSC vs. SSC) cluster tightly as a function of loaded dye concentration, with both instruments clearly discriminating 1-100 *μ*g/mL labeled BSA from background **(Fig. 3)**. The Aria II cytometer was capable of detecting < 0.1 *μ*g/mL dye (5 orders of magnitude) while the lower limit of detection for the SH800 was ~ 0.1 *μ*g/mL (4 orders of magnitude). Compared to the SH800, the Aria II cytometer and similar instruments use significantly higher-powered individual lasers; their low-range performance is equally superior on cells. Peak-to-peak separation respective to each dye series is wider on the Aria II (**Fig. 3B,C,E,F**); this increased signal discrimination is likely due to the gel-coupled collection optics and higher-powered lasers used in the Aria II (**Fig. 3B,C,E,F**). By contrast, the SH800 cytometer has a single fiber optic without gel-coupling, which reduces light collection efficiency. The SH800 also uses lower-powered lasers which reduce fluorochrome emissions and, therefore, sensitivity.

**Fig. 3.**
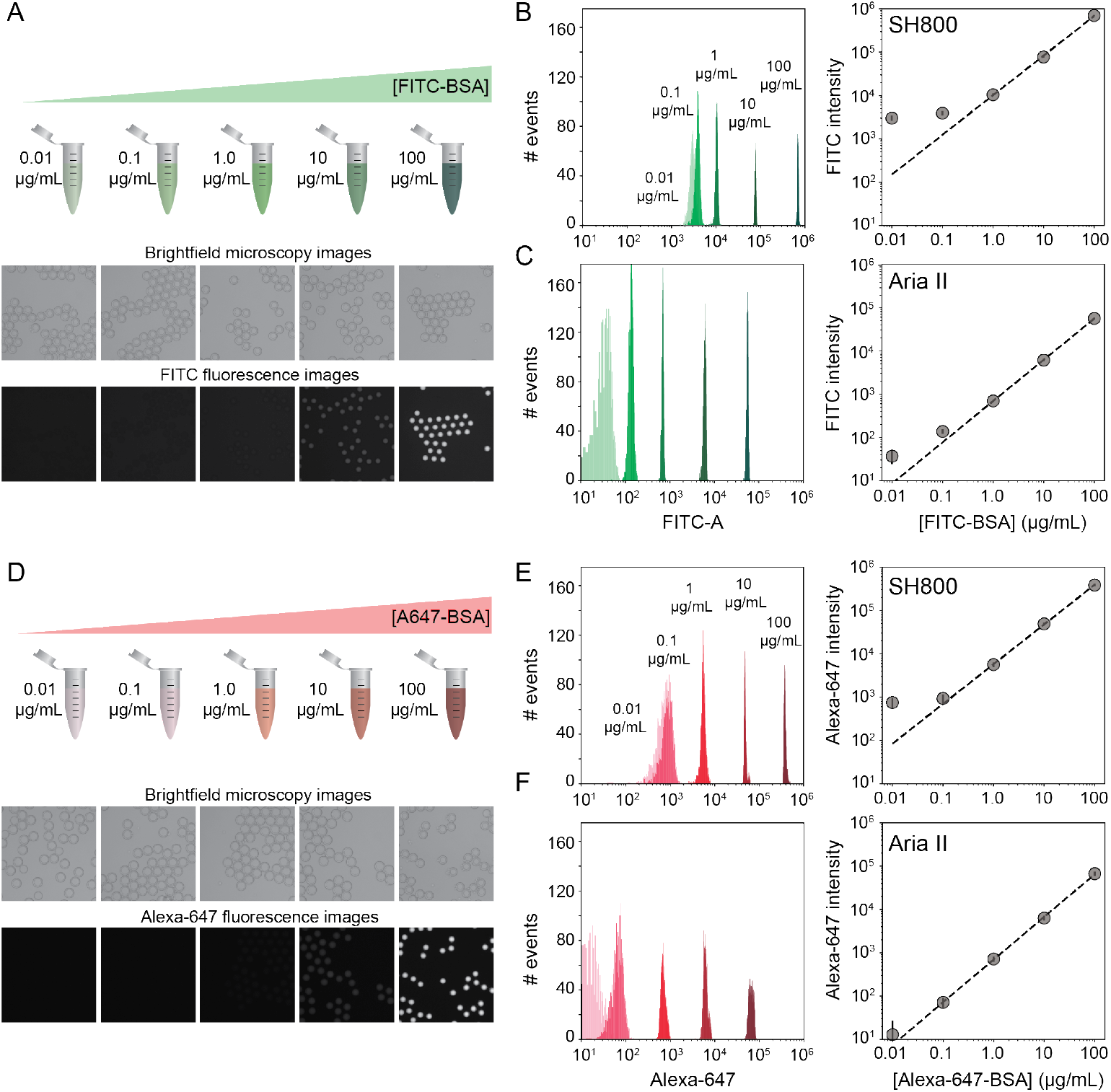
Dynamic range and limit of detection of DEs via sdDE-FACS. **(A)** Schematic and brightfield and fluorescence images of DE droplets containing multiple concentrations of FITC-BSA. **(B,C)** Histograms (left) and relationship between measured intensities and concentration for DE droplets measured on the SONY (B) and Aria (C) sorters (2,500 events/condition). **(D)** Schematic and bright field and fluorescence images of DE droplets containing multiple concentrations of Alexa647-BSA. **(E,F)** Histograms (left) and relationship between measured intensities and concentration for DE droplets measured on the SH800 (E) and Aria II (F) sorters (2,500 events/condition).

The number of photons emitted by individual cells stained with common dyes is equivalent to the 1-10 *μ*g/mL range of these calibration series (as determined by microscopy), establishing that sdDE-FACS should be compatible with typical single-cell assays. Signal variance of labelled droplet populations (peak-width of each population across the calibration series) reported here is significantly narrower than previously reported for double emulsion flow cytometry ^46^ (0.1 decade peak-width compared to ~1 decade peak-width for 0.1 *μ*g/mL FITC-BSA in the Aria II; ~10-fold improvement), allowing for more precise quantification of both high- and low-range signals.

### 2.6 Target enrichment for DE-FACS sorting

After detection, accurate sorting of individual DE droplets requires that DE droplets of interest are encapsulated within “FACS droplets” that are charged for sorting (*e.g.,* that the sample stream stably breaks into individual droplets with registration maintained so that the correct target is enriched in the destination well)(**Fig. 2B**). This registration depends on a calibrated droplet delay, which sets the delay time between when a DE droplet passes through the laser excitation and when charge is applied to target droplets after they leave the nozzle and reach the stable breakoff point ^45^. To determine the drop delay, both instruments use small (<10 *μ*m), non-deformable calibration beads during instrument setup (*e.g.* AccuDrop beads). If a droplet delay is correct for a particle sample, sort efficiencies for that particle will be maximally efficient at that delay, without compromise to target specificity in post-sort enrichment (*e.g.*, the correct particle population will be targeted, deflected, and recovered post-sort) ^52^. Sorting efficiency is calculated as a percentage of the number of recovered particles (in this case, DEs) as compared to the number of desired particles targeted for sorting; typical single-cell FACS sort efficiencies range from 50-90% by cellular type and size ^36,43,51–53^. For the SH800, the bead-calibrated droplet delay was found to be optimal for DE post-sort recovery efficiency. By contrast, sorting efficiency on the Aria II was optimal at droplet delays significantly outside the Accudrop values (**Fig. S7**); droplet delay must be determined empirically for each droplet size on the Aria II. These differential effects of droplet delay are likely due to different flow metering and acoustic droplet mechanisms between the two instruments (**Table 2**); in the case of the Aria II, calibration beads may not accurately represent large-cell or droplet dynamics. For the Aria II, an additional calibration step to manually adjust droplet delay for DEs using a control DE population is required for accurate target enrichment and post-sort recovery.

To demonstrate the utility of sdDE-FACS for rare population enrichment and validate empirically-determined droplet delay times on the Aria II, we attempted to enrich for a population of FITC-BSA-loaded DE droplets present at 20.4% of a parent population of blank DE droplets using both sorters (**Fig. 4**). Bright-field and fluorescence images of pre-sort droplet populations confirmed that FITC-BSA droplets were present at the target mixed abundance (**Fig. 4A,B**) and side *vs.* forward scatter profiles on the SH800 and Aria II showed a distinct cluster of DE droplets for the mixed population (**Fig. 4C,D**).

**Fig. 4.**
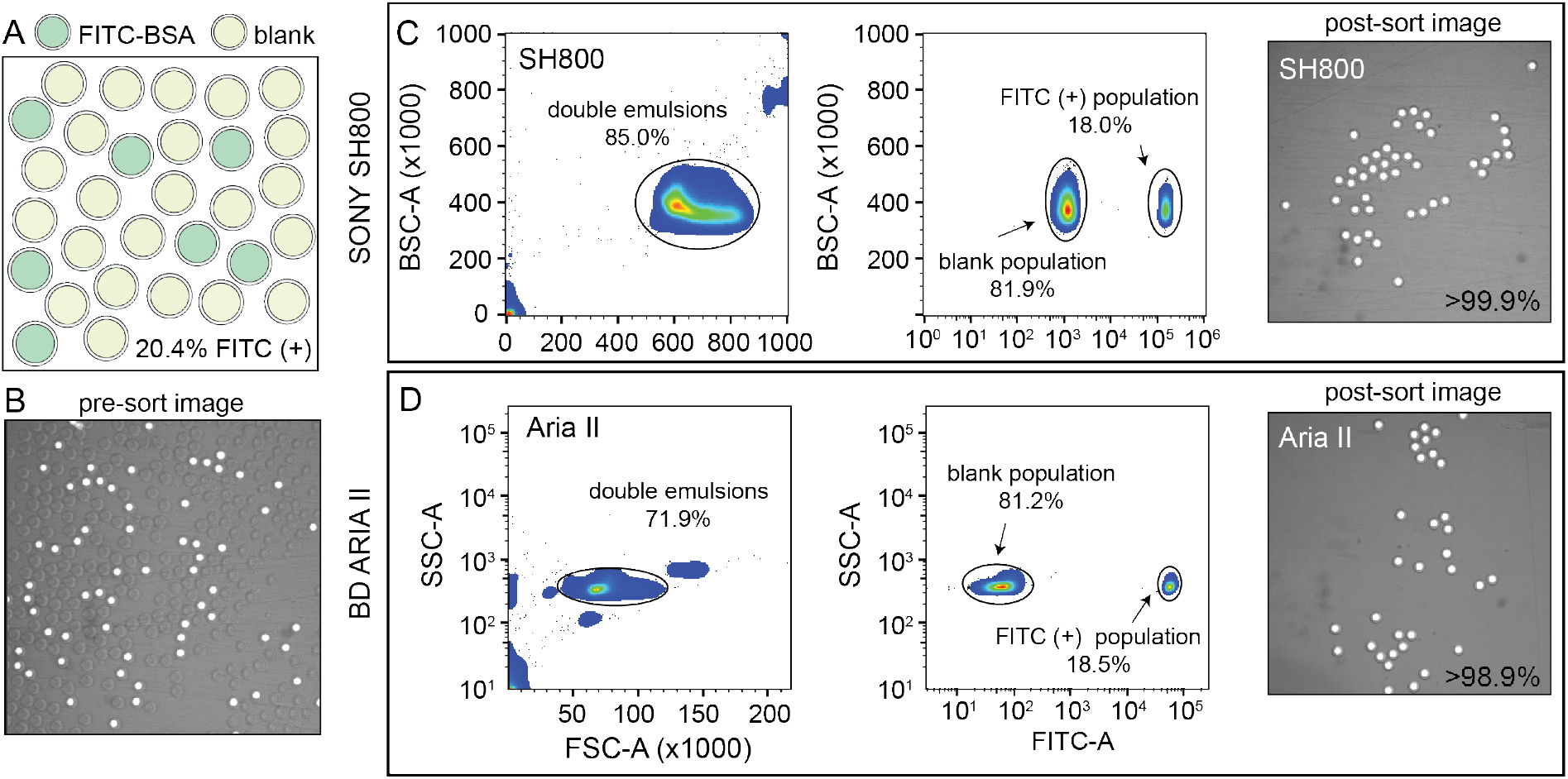
Rare population target enrichment via sdDE-FACS. **(A, B)** Schematic and microscopy image of pre-sort DE droplet populations containing 20.4% FITC-positive droplets as determined by manual count (62/303 droplets positive). **(C)** SH800 FACS gates and post-sort image of 100 droplet well with associated target enrichment sensitivity. **(D)** Aria II FACS gates and post-sort image of 100 droplet well with associated target enrichment sensitivity. Parental FACS gate shows 10,000 events/condition, randomly sampled, for both sorters.

Investigation of measured FITC intensities revealed clearly separated populations of blank and FITC-positive droplets consistent with expectations, with a good parental population estimates on both instruments (18.0% and 18.5% for the SH800 and Aria II, respectively) (**Fig. 4C,D**). Results are consistent across different flow cell geometries (square flow cell replicate, **Fig. S8**). Post-sort, both the Aria II and SH800 showed near-perfect enrichment of intact target DE droplets (>98.9% and >99.9%, respectively, with 0 false positives observed over multiple fields of view for the SH800) (**Fig. 4C,D**). These results confirm the ability to selectively enrich post-sort for “rare” droplet populations with high target specificity via sdDE-FACS.

### 2.7 Single droplet sorting using DE-FACS

Accurately linking genotype to phenotype for *individual* selected variants at high-throughput requires that droplet recovery be maximally sensitive in selecting for the correct variant from a mixed population. However, to enable high-throughput single-cell droplet applications, such as single cell sequencing from droplet microreactors, droplet sorting must also be maximally efficient (*e.g.,* as many wells as possible are occupied by a single DE droplet). Further, droplets must also remain intact during sorting to prevent cross-contamination of material between wells. To quantify sorting efficiency, we generated populations of DE droplets and attempted to sort 100, 10, or 1 droplets into alternating wells of a 96-well destination plate containing fluid osmotically matched to the DE droplet core (**Fig. 5**). Empty wells systematically interspersed between destination wells enabled testing for spray-based DE droplet cross-contamination. After FACS droplet deposition, we imaged all wells and manually counted the number of recovered droplets **(Fig. 5b)**. Across droplet populations and for each plate, wells designed to contain 100 or 10 droplets contained on range of [64.4 - 83.6] and [5.9-8.4] average droplets (n=12-36 wells per plate per 100- or 10- droplet set points), respectively, for the Aria II. For the SH800, wells designed to contain 100 or 10 droplets contained on range of [46.2 - 69.9] and [4.9-7.1] average droplets (n=12-36 wells per plate per 100- or 10- droplet set points), across droplet populations, for an estimated achievable droplet recovery rate of 60-80% for both instruments (**Fig. 5C; Table S3**), dependent on droplet condition. Droplet size and oil shell thickness had minor effects on sort efficiency after adjusted droplet delay (**Table S3, Figure S7**). These quantitative estimates represent the first reported plate statistics by for DE recovery. Imaged droplets remained intact **(Fig. S9)**, and out of 193 total wells, only 5 negative control wells designated to contain 0 droplets were observed to contain a droplet.

**Fig. 5.**
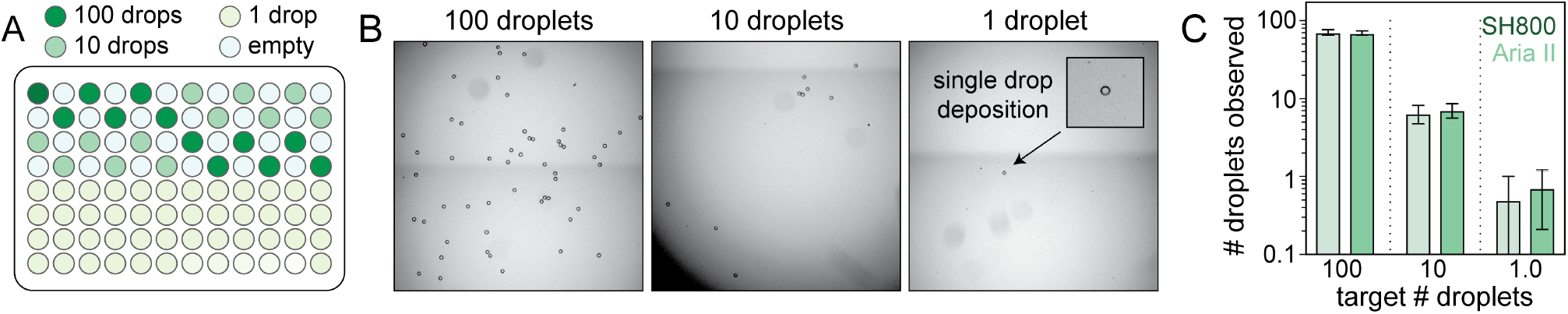
Plate-sorting statistics for DE droplets using sdDE-FACS workflow. **(A)** Schematic showing an target FACS droplet deposition in a 96-well plate (exemplary plate format; some plate formats contain greater or fewer 100, 10, or 1 droplet target wells, as noted). **(B)** Brightfield images of individual wells within a deep-well optical 96-well after sorting to deposit 100, 10, or 1 FACS droplets (each containing a single DE droplet). **(C)** 96-well plate sorting statistics for a representative DE droplet population. Mean and SD error bars shown. Means by set point (Aria II, SH800): 100 droplets (71.2, 69.9; n=11-12 wells), 10 droplets (6.1, 7.1; n=36 wells), 1 droplet (0.5, 0.71; n=24 wells). Additional plate statistics are available in **Table S3**.

Most importantly, single DE droplets can be reliably sorted and recovered via sdDE-FACS. Wells designated to receive a single DE followed a bimodal occupancy distribution where wells either contained a single deposited droplet or no droplet at all (**Fig. 5C, 6B**). Single droplet recovery efficiencies were typically ~ 70% ([0.5-0.83] droplets, n= 36-48 single droplet wells per plate), with the highest sort efficency (83%) observed on the Aria II **(Table S3)**. Single droplet plate counts (**Fig. 5C, Table S3**) are consistent with prior recovery efficiency estimates for single-cell deposition via FACS ^51–53^, suggesting sdDE-FACS has reached instrument limits.

**Fig. 6.**
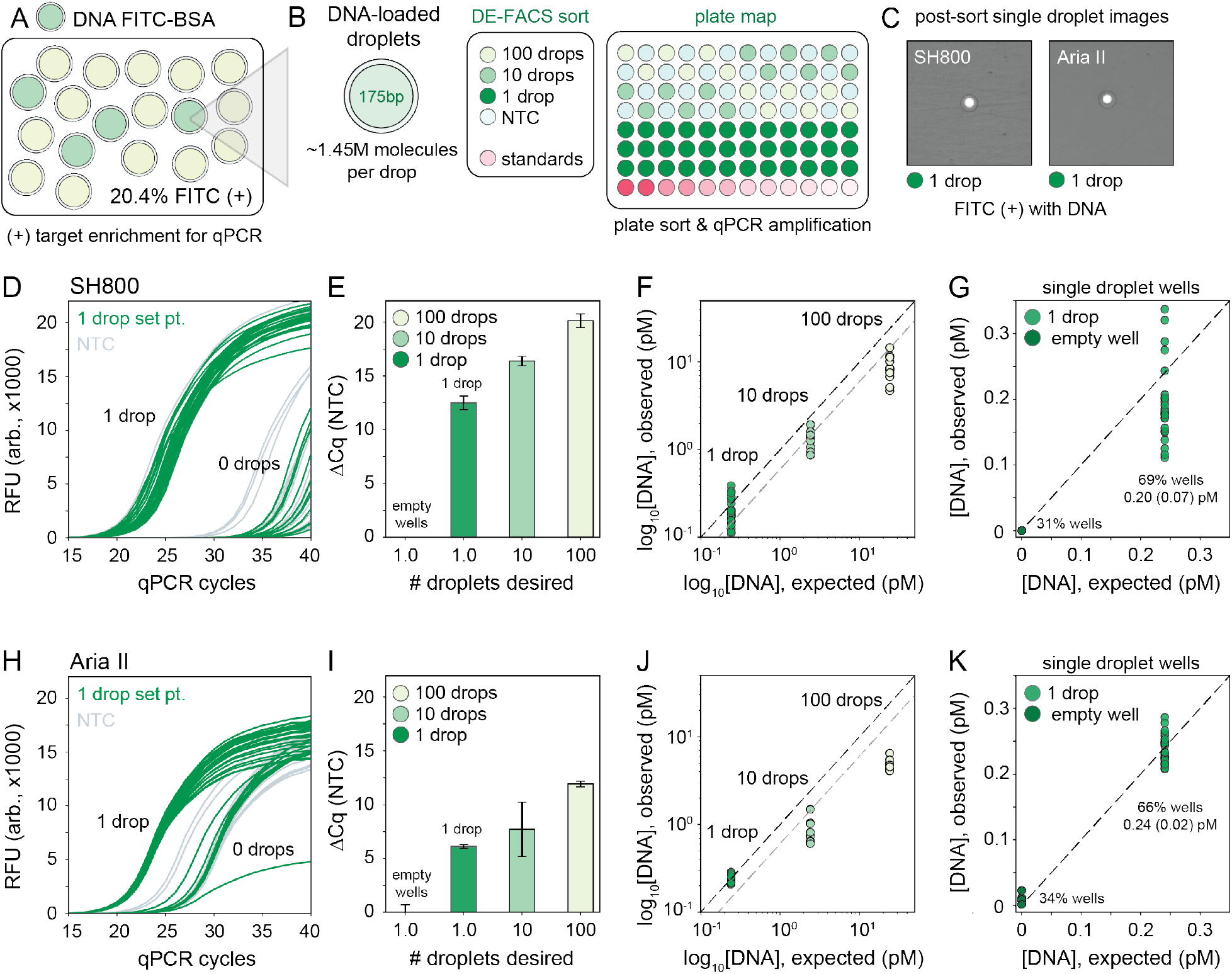
Downstream nucleic acid recovery and qPCR processing of DE droplets with sdDE-FACS. **(A)** Mixed populations containing DNA-loaded DEs labelled with FITC-BSA in a pool of blank droplets. **(B)** Plate map schematic for the qPCR assay (10 *μ*L reactions). **(C)** Optical confirmation of post-sort enrichment for the SH800 and Aria II on single droplet wells. **(D,H)** Raw qPCR traces for single droplet wells (n=36 wells/plate) for the SH800 and Aria II, respectively. **(E,I)** Cycling thresholds (NTC subtracted) for all droplet-designated wells for the SH800 and Aria II. **(E,J)** Absolute quantification using in-plate DNA standards for 100, 10, and 1 droplet per well set points with further analysis of single droplet designated wells, **(G,K)**, as clustered by bimodal sorting statistics for the SH800 and Aria II, respectively. Mean (SD) recovered DNA concentrations are indicated; expected concentration is 0.24 pM per well. Each point represents an individual well measurement.

### 2.8 Genotyping of double emulsions using DE-FACS

Successfully linking *genotype* to *phenotype* for individual variants via downstream nucleic acid interrogation assays (*e.g.* qPCR or next-generation sequencing) is a critical application of single cell FACS ^40,41,44^. Enabling similar capabilities for single DE droplet microreactors requires that DNA be recoverable from each droplet post-sort without loss of material or cross-contamination from breakage during aerosolization or sorting.

To quantify nucleic acid recovery and sensitively detect cross-contamination, we generated a library of DE droplets (27.8 *μ*m total diameter, 16.6 *μ*m core) each containing ~1.45 million molecules of a small 175-bp DNA fragment (a portion of the coding sequence for the GAPDH housekeeping gene) and 10 *μ*g/mL FITC-BSA. DNA-containing DEs were mixed into a blank DE population of the same droplet size (**Fig. 6A**). Using the Aria II and SH800 cytometers, we then sorted 100, 10, or 1 DE droplets, targeted by FITC fluoresence, into alternating wells of a destination plate with in-plate DNA standards and systematically interspersed empty wells (**Fig. 6B**). Target droplet isolation was confirmed optically per each sort (**Fig. 6C**). DE droplets were lysed by depositing FACS-generated aerosolized droplets (containing single target DE droplets) into dry wells of a multiwell plate and allowed changing interfacial tension to drive DE droplet lysis within 30 seconds **(Fig. S10)**. This approach outperformed previously published methods that rely on bath sonication or chemical breakage of the emulsion (a known inhibitor of PCR) ^13,59^. To sensitively quantify DNA recovery, we then performed plate-based qPCR using GAPDH-specific primers; serial DNA standards were included for calibration **(Fig. 6B)**.

Across both instruments, cycle thresholds (Cq) for wells designed to contain 100 or 10 droplets cluster tightly within each group, with lower spread in across all droplet set points observed on the SH800 **(Fig. 6E,I)**. Absolute quantification of the amount of DNA recovered from each well using in-assay standards demonstrates full recovery of all DNA from single DE droplets (**Fig. 6F, J**). Between 1 and 10 droplet-containing wells an expected near 10-fold drop in recovered DNA is observed, as modulated by expected 60-70% sort recovery efficiency (**Fig. 6F, J**, blank line indicates 100% sort efficency, grey line indicates projected 60% sort efficency). Notably, the SH800 more accurately replicates expected DNA recovery by sort efficiency. Between the 100 and 10 droplet samples, reduced recovery of nucleic acids beyond 60% sort inefficencies is likely due to decreased droplet lysis using the dry plate technique **(Fig. S10)**; 30-60s of drying time is likely insufficent to fully evaporate 100 “FACS droplets,” each 6.9 nL in volume (total volume: 0.69 *μ*L). Octanol extraction (using PFO) or similar bulk-based droplet lysis techniques may perform better in 100+ droplet recovery regimes ^13,59^. Across both instruments and at all desired droplet set points, DNA concentrations are within size variation error or below 100% recovery expectations, indicating minimal droplet-breakage and correspondingly minimal DNA cross-contamination from global droplet lysis in the FACS stream.

Of critical importance, single droplet nucleic acid recovery via sdDE-FACS reliably reproduces expected DNA concentrations at high sort efficiencies (69%, SH800; 66% Aria II, single droplet isolation, **Fig. 6G,K**). qPCR traces for wells targeted to contain a single DE droplet show bimodal clustering consistent with optically-derived plate sorting statistics, with >60% of single droplet wells observed in a clearly discernable set in the raw qPCR traces (remaining single droplet wells cluster with no template negative control wells suggesting 0 droplet occupancy, **Fig. 6D,H**). Further investigation of single droplet wells with 1 or 0 droplet occupancy reveals excellent concordance with expected DNA concentration in wells containing a single droplet (0.20 ± 0.07 pM, SH800; 0.24 ± 0.02 pM, Aria II; 0.24 pM, expected). Results are consistent across replicates and additional populations **(Fig. S11)**. These data indicate successful, complete recovery of nucleic acids via downstream qPCR in isolated single DEs via sdDE-FACS.

## Conclusions

Here, we have demonstrated a high-throughput DE droplet sorting and recovery strategy (sdDE-FACS) capable of quantitatively phenotyping, sorting, and recovering nucleic acids from droplet subpopulations or individual droplets using standard FACS cytometry. The ability to robustly analyze and sort droplet populations via FACS using the optimized settings described in this work increases droplet sorting throughput by approximately an order of magnitude relative to typical FADS-based sorting and achieves high-throughput plate-based single droplet isolation ^19,22,24^. Further, droplet sorting with FACS unlocks unparalleled signal quantification and analysis parallelization capabilities of multi-color flow cytometry currently inaccesible to FADS-based sorters ^2,29,33,34^. As microfluidic droplet generation is a high-throughput process (1-100 M droplets per reaction; 10-30 kHz production rates), this increased plate sorting throughput, signal discrimination, and ease-of-access allows realization of the full potential of droplet microfluidics for high-throughput screening of rare variants.

sdDE-FACS significantly lowers the barrier to entry for new droplet sorting assays, with no need for specialized equipment beyond that typically available at many universities and companies. The syringe pumps and setup used for emulsion generation are inexpensive and widely available and we demonstrate that sdDE-FACS is compatible with 2 industry-standard sorting instruments. To facilitate broad adoption across labs, we provide detailed information about optimized instrument settings (laser settings, flow rates, and drop delays) and droplet formulations (surfactant, oil, and buffer mixtures).

Beyond enhancing the throughput and accessibility of droplet sorting, sdDE-FACS represents a critical first step towards realizing a broad suite of novel single-cell analysis assays. As one example, difficulties associated with buffer exchange in droplet microfluidics have limited the number of tandem assays that can be performed on the same single cell, requiring that researchers first identify a single buffer moderately compatible with both assays by tedious trial and error. sdDE-FACS enables a broad range of new “multi-omic” assays by making it possible to perform a first assay within a microfluidic droplet and then transfer this droplet to a well of a destination plate containing a second buffer ^39,44,60–62^. As the destination well contains a significantly higher volume than the droplet (~10,000X), near complete buffer exchange between reactions is possible enabling optimal performance of both reactions. This scheme also preserves the ability to link any quantitative fluorescent phenotype to cellular measurements, facilitating analyses of single-cell intracellular proteins, secreted proteins, labelled nucleic acids, or cellular activity (*e.g.* pH change or treatment response). If combined with existing droplet single cell genomic and transcriptomic measurements ^3,12,13,59^, cellular phenotype and genotype could be directly linked within each microreactor. The strigent single droplet isolation as enabled by sdDE-FACS allows current assays (*e.g.* high throughput enzyme and functional product screening ^2,5,63^) to achieve superior quantitative capabilities (*e.g.*, improved reaction product discrimination), improved ease-of-use (*e.g.* no sorter clogging), and efficient subpopulation enrichment and rare variant isolation.

sdDE-FACS presents an optimized pipeline for DE droplet generation, phenotyping, and sorting down to single droplet isolation compatible with FACS. Similar to the trajectory of single cell FACS, we hope this technique will be broadly applicable to novel, highly quantitative droplet assays that link variant phenotype to genotype.

## Materials and Methods

### Double emulsion device fabrication

Monodisperse DEs were generated using a one-step co-axial dual flow focusing device with flow filters and a flow resistor, similar in design to previous reports ^56,57^ (**Fig. S1**). Microfluidic Si wafer master molds were constructed using standard photolithography techniques with a 15 *μ*m relief height for the first flow focuser (to generate water-in-oil emulsions) and a 40 *μ*m relief height for the second flow focuser junction (to generate water-in-oil-in-water emulsions) using 2-layer SU8 2015 deposition prior to a development step. Poly(dimethlylsiloxane) (PDMS) microfluidic devices were fabricated from the master molds using soft lithography at a 1:5 elastomer base: crosslinker ratio. Post-bake, droplet generation devices were hole punched using a 1 mm biopsy punch (PicoPunch) and monolithically bonded to a blank 1:10 PDMS slab (5 cm in height). Devices were baked for 48 hours, with longer baking times improving hydrophobicity of the resultant droplet generation devices. Immediately prior to generating double emulsions, the device outlet path was selectively O2 plasma treated for 4.5 min at 150W plasma (Fempto, Diener) by taping device inlets. This process allowed for the outer flow focuser of the device to switch to hydrophilic wettability while retaining hydrophobicity at the first flow focusing junction ^57^.

### Double emulsion generation

Double emulsions were generated using 3 syringe pumps (PicoPump Elite, Harvard Apparatus) for the inner, oil, and carrier fluids. The inner phase for the aqueous droplet core was composed of Tween-20 (Sigma) in PBS (Invitrogen), with additional reagents (*e.g.* FITC-BSA, Invitrogen) as indicated in (**Table 1**). BSA (0.5-2%) can be optionally substituted for Tween-20 (0.1-1%) in the droplet core to no adverse effect. The oil phase was composed of HFE7500 (Sigma) and Ionic PEG-Kyrtox (FSH, Miller-Stephenson). The carrier phase contained Tween-20 (Sigma) and Pluronix F68 (Kolliphor 188, Sigma) in PBS. Each phase was loaded into syringes (PlastiPak, BD; Hamilton, Sigma, see Extended Methods), and connected to the device via PE/2 tubing (Scientific Commodities). Typical flow rates were 275:75:2500 (oil: inner core: outer aqueous sheath) *μ*L/hr. Droplet generation was monitored and recorded via a sterescope (Amscope) and high-speed CMOS camera (ASI 174MM, ZWO) (**Fig. S1**).

### Preparation of double emulsions and instruments for FACS

Prior to FACS sorting, double emulsions were diluted 1:5 in FACS diluent buffer in a 12 × 75 mm round bottom FACS tube (BD Biosciences). For a typical run, 100 *μ*L of double emulsion droplets were removed from the droplet pellet (containing high surfactant outer mix) and adding them to 500 *μ*L of FACS diluent. Droplets were gently resuspended before analysis. See Extended Methods in *Supplemental Information* for further discussion. Both instruments were thresholded on forward scatter, FSC, a sizing parameter, at extremely low values since DE droplets are large compared to typical cells (**Table 1**). Sort gates were widely permissive to show droplet purity (including sample free oil and dust, if present) compared to background. Thresholding is indicated in figure legends, if applicable. Event rates were capped below 300 events/s during sorting and 1000 events/s during analysis-only runs by modulating flow rate or flow pressure; the initial appearance of DE droplets for the Sony SH800 was typically delayed 100-200s (see Extended Methods, **Fig. S3**). All post-processing analysis was completed in FlowJo v10.5.3 (FlowJo) and using custom Python scripts.

### FACS Analysis on Aria II (BD)

DEs were loaded and analyzed on the FACS Aria II (BD) using a 1.5 ND neutral density filter in the optical path to visualize the droplet population and a 130 *μ*m nozzle for sorting. Droplets were first gated on FSC and SSC profile, followed by singlet gating using FSC-H and FSC-A and subsequent gating on APC, FITC or DAPI fluorescence, as indicated. All flow and thresholding parameters are reported in **Table 3**. Sorts were completed with single cell purity mode.

### FACS Analysis on Aria II (BD)

DEs were loaded and analyzed on the FACS SH800 (Sony) using a standard 408 nm laser configuration and a 130 *μ*m microfluidic chip for sorting. Droplets were first gated on FSC and SSC profile, followed by singlet gating using FSC-H and FSC-A and subsequent gating on APC, FITC or DAPI fluorescence, as indicated. All flow and thresholding parameters are reported in **Table 3**. Sorts were completed with single cell purity mode.

### Plate sorting of double emulsions

Plate sorting was conducted using 96-well optical plates (Fisher Scientific) or qPCR plates (Biorad) on the Aria II and SH800 using associated 96-well plate gantries for each instrument. Prior to sorting, 100 *μ*L of osmotically-balanced outer phase buffer was loaded into each well. Optimal drop delay was calculated for the Aria II instrument by using a blank droplet population, run the same day as the sample of interest. A protocol is available in Extended Methods. Briefly, blank droplets were sorted at set point of 50 droplets per well after Accudrop calibration and laser compensation, with each well corresponding to a different droplet delay setting (manually input) from −2.5 to +2.5 delay units in increments of 0.25 delay units from the Accudrop automatic droplet delay **Fig. S7**). Droplets were manually counted using a low-cost benchtop stereoscope (Amscope) to decide on the highest efficiency drop delay per the population; the process takes ~ 5-10 minutes and is a recommended step in calibration. Plate statistics were determined by 96-well optical images (EVOS microscope, 4X Objective, Life Technologies) and manual counting. High-resolution droplet imaging used for size analysis and visualizaiton was was conducted using a Ti Eclipse microscope (Nikon) and sCMOS camera (Zyla 4.2, Andor) at 10X (16-bit, low-noise) with Brightfield Dichroic and eGFP filter sets (Semrock).

### Nucleic acid recovery and droplet genotyping

qPCR assays were conducted using the iTaq SYBR I qPCR master mix (BioRad) on a CX96 qPCR instrument (Biorad). Primers and fragment sequences are available in **Table S4**. Droplets were lysed into a dry qPCR plate (Biorad) (see Extended methods, **Fig. S10**) for 1 min post-sort. Immediately after, 10 *μ*L reaction mix **Table S5** was added per well as shown in (**Fig. 6A**) and in-assay standards were added subsequently in the remaining row. The reaction was thermocycled according to the following program: 2 min 95C, [95C 0:05s, 60s 0:30s]×50. A melt curve with 2C increments from 65C - 95C was performed after each run to distinguish on-target amplification from primer-dimer amplification.

### Droplet Size Characterization

Droplets were characterized via a custom MATLAB script available via our Open Science Framework repository; methods are outlined in *Supplemental Information*.

### Open Science Framework Repository

An OSF Repository is available for this project containing data, images, and associated software for this method, and is located at **DOI: 10.17605/OSF.IO/3AU4V**.

## Author Contributions

K.K.B, S.K., and P.M.F. conceived and designed the study. K.K.B. designed the methdology. K.K.B. and C.C.C. performed experiments with assistance from B.C. and S.G.C. G.K. and K.B. contributed software. S.K. and B.C. contributed resources. K.B., C.C.C., B.C., S.G.C., and G.K. conducted formal analysis, data curation, and validation, with input from L.N. and P.M.F. K.K.B., L.N., and P.M.F. wrote the manuscript, with input from all authors.

## Acknowledgements

We acknowledge Dr. David Sukovich and Dr. Adam Abate for early discussions related to optimizing droplet FACS, demonstration of their flow cytometry workflow, and for the use of their device design. We also acknowledge Dr. David Parks, Dr. Lee Herzenberg, Dr. Aaron Cantor, and the Stanford Shared FACS facility for project guidance, advice, and technical support. We thank the Fordyce Laboratory and the Wang Laboratory, in particular Margarita Khariton, for their critical feedback and comments on the review of this manuscript. We additionally thank the Stanford FACS Core for use of their instruments and facilities as well as the Stanford Protein and Nucleic Acid (PAN) facility, in particular Dr. Mike Eckhart, for use of the Life Technologies EVOS microscope. K.K.B. acknowledges funding as Chem-H CBI fellow, an NSF GFRP fellow, and a Siebel Scholar. This work was supported by NIH grant 1DP2GM123641, and P.M.F. is a Chan Zuckerberg Biohub Investigator.

## Supplemental Information

## Extended Materials and Methods

### 1. Flow Cytometry Preparation of Double Emulsions

Prior to FACS, DEs were diluted 1:5 in FACS diluent buffer (1% Tween-20) in a 12 × 75 round bottom FACS tube (BD). For a typical run, 100 μL of DEs from the DE pellet at the bottom of the collection vessel (***e.g.*** 1.5 mL Eppendorf) were gently aspirated with a P200 pipette prior to dilution. All DEs were osmotically matched to their suspension media (the outer aqueous sheath high-surfactant mix containing 2% Pluronix F68 (Sigma), **Table 1**) to prevent DE expansion or shrinkage. To process smaller absolute amounts of DE droplets (<100 μL pellets), we recommend supplementing the remaining volume with the outer aqueous sheath high-surfactant mix before dilution in the FACS diluent buffer to increase droplet stabilization during sample loading and FACS injection into the sample line.

#### Suggestions for Preparation of Double Emulsions for FACS

- Always use at least 50 μL of droplets from the droplet pellet, if possible, as too few droplets will require sample pausing, manual agitation, and reloading.
- Before loading droplets into the FACS machine, gently swirl or flick the FACS 12 × 75 mm tube, but <u>never vortex.</u> Droplet should be gently resuspended from the white pellet to create a cloudy mixture but vortexing may induce too much shear and result in droplet breakage.
- Manual resuspension may be required during the run, especially in the SH800. If event rate drops below 50 eps after initial gate structuring, pause and resuspend the droplets by gently swirling the sample. Sample line re-positioning to the bottom of the FACS tube can also significantly decrease the need for resuspension.
- Density matching agents (***e.g.*** OptiPrep, xanthan gum, or PEG) may be used but can change typical FSC vs. SSC profiles; the SH800 does not have optionality for an added ND filter and droplet populations may not appear on-scale using these additives. Manual resuspension performs more robustly and reliably for high sort recovery in our hands and is strongly recommended.
- Optically dense samples (e.g., greater than 150 μL of droplets loaded per 500 uL) may result in decreased droplet singlet rates and droplet breakage. Droplets are deformable and can pack in sample loops and nozzles; too many droplets run at once can collide and result in breakage. To process large numbers of droplets, we recommend pausing and continuing to add sample during the run rather beginning with dense samples. The time required for droplet addition is minimal.

### 2. Droplet Delay Large-Particle Manual Calibration

Before each run, droplet delay times were calibrated using instrument software for the SH800 (Sony) and Aria II (BD) using flow calibration beads (Automatic Setup Beads, Sony; Accudrop beads, BD), typical of traditional FACS workflows for cellular analysis. Bead-calibrated drop delay values and starting droplet spacing, droplet profile, and drop-drive frequency were recorded as set points before manual droplet delay calibration. Manual adjustment of the droplet delay had minimal effects on droplet recovery efficiencies for the SH800; intital droplet delay values were used for all SH800 sorts. For the Aria II, we observed a significant effect on sorting efficiency upon adjustment of the droplet delay (see **Fig. S7**, and Extended Notes). A manual droplet delay adjustment using control DE populations prior to each run was performed on the Aria II. Confirmatory droplet delay calibrations can be confirmed on either instrument, as desired.

#### Protocol for Manual Droplet Delay Adjustment

**Run time:** 5-7 min

1. Generate droplets for manual droplet delay adjustment by either (1) setting aside a small split-fraction of your droplet sample (dedicated for calibration-only) or (2) generate a small (^~^20 μL pellet, 2-4 min generation time) control droplet population using the same flow rates (recommended for rare or precious samples). Calibration droplets must have a similar total volume and oil shell size, as these parameters influence the resultant DE-optimized droplet delay.
2. Calibrate the cytometer using standard flow cytometry beads and note the drop delay set point.
3. Load the chosen calibration droplet population and gate according to FSC vs. SSC morphology.
4. Load an optical 96-well plate into the plate loader. Each well in 1 row should contain 100 μL of outer aqueous sheath buffer to stabilize the sorted droplets.
5. Using the instrument software, manually adjust the drop delay units on the droplet profile from the set point by −1.5 to +1.5 in 0.25 – 0.5 unit increments, as in **Fig. S7**.

5.a. For each increment, sort 50 droplets with single cell purity into a well of the 96-well plate. Note the well location and droplet delay per each sort.
5.b. Wait 10 s during active event collection at the new droplet before proceeding to the next sort. This stabilizes target selection to ensure droplets of the right drop delay enter the well.
5.c. If FACS droplet profiles, droplet-drive settings, or spacing become unstable, wait for stabilization or adjust the parameters until stabilization is regained. Droplet delay profiles must be collected under stable FACS droplet breakoff.
6. When the droplet delay calibration sort has been completed, manually count each well using a benchtop stereoscope (Amscope). This step should take 2-3 min total. Automated imaging is possible (e.g., using an EVOS plate microscope, Life Technologies) but is not necessary.
7. Select the adjusted droplet delay profile with the maximal number of droplets collected in the sweep. Successful calibrations should achieve 50 – 90% sort recovery efficiency; set point recovery efficiencies are predict later sort statistics for the population. Proceed with all further sorting and analysis using the new droplet delay.

We have found manually-calibrated drop delays (as expressed relative to the Accudrop-calibration value, ***e.g.*** +0.5 d.d. units) are extensible to DEs of the same droplet size and chemistry.

### 3. Droplet Size Analysis

Droplets were imaged for size characterization on a Nikon Ti Eclipse microscope at 10X under brightfield and a fluorescence channel according to a reference dye in each droplet. In this work, FITC was used as the reference dye. Multiple images were analysed per droplet population (typically, 5 – 20 images per condition, 50 – 500 droplets analysed total per condition). An automated pipeline was developed in MATLAB to extract radial and volumetric information about the droplets. First, a central ROI of 500 x 500 pixels was extracted from each droplet image. In the FITC channel image, center coordinates and radii of all inner cores were obtained using the circular Hough transform method (imfindcircles). In the brightfield channel image, each droplet was isolated in a square ROI centered at the x- and y-coordinates of the extracted droplet centroid, as determined from the fluorescence image, with each side twice the length of the inner core’s radius. The droplet ROI was contrast-enhanced and coverted to grayscale through contrast-limited adaptive histogram equalization (adapthisteq). An image profile of pixel intensities was taken along the horizontal and vertical center lines of the ROI (improfile). Given that the outer boundary of a droplet has low pixel intensity relative to its surroundings, intensity and positional information of local minima (identified using peakdet) in the image profiles were extracted. By direct comparison of the coordinates of the first and last minima in the horizontal and vertical profiles, the two positions closest to the droplet’s center were selected to demarcate the outer shell’s boundaries. If the two minima originated from different image profiles (i.e. one from the horizontal profile, and the other from the vertical profile), positions were standardized by converting the last minimum to the minimum closest to its original position on the other image profile. The droplet diameter was reported as the difference between the two extracted minima. To correct overlapping droplet boundaries, the difference between the sum of the overlapping droplets’ radii and the Euclidean distance between their centers was subtracted from each droplet’s radius. Droplets whose outer radii were found to be smaller than their inner radii were removed from consideration. Through visual inspection, droplets were excluded from analysis during outer shell boundary detection if a local minima could not be identified for a given droplet due to close droplet packing (typical rejection rate: 10%, random sampling).

Droplet sizes were manually quantified in samples lacking a reference dye (e.g., PBS Blank droplets, **Fig. 2**). Brightfield images were opened in ImageJ, and circular regions of interest were manually drawn around each inner core and outer shell visible in an image. For each population, 50 identified droplets were randomly sampled and reported for size characterization; ROI coordinates and masks were saved and recorded for each population.

### 4. Droplet Taqman PCR and Thermocycling

To that sdDE-FACS is compatible with thermocycling reactions, we performed in-droplet genomic Taqman PCR in double emulsions as described in Sukovich et al., 2017^1^ and subsequently sorted Taqman-positive droplets using the sdDE-FACS technique. Lambda DNA (NEB) and Phi X174 DNA (NEB) were diluted in qPCR-grade water (Invitrogen) prior to use. A PCR reaction was assembled as follows: 25 μL Platinum Multiplex PCR Master Mix (Thermo Fischer), 2.5 μL Taqman probe (5 μM), 0.5 μL forward primer (100 μM), 0.5 μL reverse primer (100 μM), 2 μL diluted DNA, 5 μL UltraPure BSA (5% stock, Ambion), and qPCR-grade water (Invitrogen) to 50 μL. A separate mix was prepared for each DNA sample. The PCR mix was loaded into a single-inlet device to generate ^~^30 μm double emulsions; flow rates were 275:75:2500 (O:I:S) μL/hr. After collection, droplets were supplemented with 2X Taq Polymerase reaction buffer (Thermo Fisher) and 1.5 mM MgCl_2_^2+^ in 1:1 dilution with the outer aqueous phase and subjected to Taqman PCR cycling using a Life Technologies thermocycler according to the following program: (86C for 2 min; 40 × 86C for 30s, 60C for 90s, 72C for 20s; 72C for 5 min). Post-cycling droplets were imaged using a Nikon TiEclipse under brightfield and FAM fluorescence. Droplets containing either lambda DNA (NEB) and Phi X174 DNA (NEB), and associated probes and primers, were mixed 1:100 in the droplet sample and analysed via FACS, with target enrichment for the positive population. Droplets were subsequently imaged post-recovery for direct comparison on the performance of the sdDE-FACS workflow to Sukovich et al., 2017^1^.

## Extended Discussion Notes

### 1. Double Emulsion Generation

We generated double emulsions were generated using 3 syringe pumps (Pico-Pump Elite, Harvard Apparatus) for the inner aqueous, oil, and outer carrier fluids as shown in **Fig. S1**. Additional syringe pumps and sample loaders (e.g., sample loops, paired syringe pumps, stopcock assemblies) can be added, as desired, for additional inner core reagents (e.g., reaction mixes, lysis buffers, cell dyes, etc.) or reagent exchange dependent on the design of the microfluidic device. Syringes can be selected for either normal-volume (1 – 5 mL; PlastiPak plastic syringes, BD) or low-volume (10 – 500 uL; Hamiliton glass syringes, Sigma) applications per reagent. We recommend the use of polyethylene tubing (PE/2, Scientific Commodities) because of its low-bind properties and biological compatibility, but Tygon, FEP, or alternative polymer tubing are also attractive options. A full description of setup components is available in **Table S1**; costs are typically between $7,000 - $10,000 for setup construction with <$10 consumables per device array (10 – 15 runs per array). Droplet generation rates are typically 1-10 kHz, with 0.6 - 6M droplets collected per each 10 min run time for 30 μm droplets. Higher or lower collection rates can be achieved with smaller or larger size droplets, respectively.

### 2. Double Emulsion Device

The device design used in this work is similar to a previously reported dual flow-focuser design2,3. In this design, single emulsions containing an inner aqueous core are first generated in an oil sheath (flow focuser #1) and subsequently enveloped in an oil shell by the second aqueous sheath (flow focuser #2), separated by a flow resistor channel of high fluidic resistance to stabilize the two flow regimes. This device allows highly uniform, reproducible droplet generation using an easy ‘one-step’ approach (e.g., all inlets are plugged into the same singular device and run simultaneously to produce resultant double emulsions). We have fabricated multiple variants of the ‘one-step’ dual-focuser device presented in this work containing a varying number of inlets for different reaction schemes, or different channel heights (***e.g.,*** a 25 μm inner flow focuser, 50 outer flow focuser device for 40 μm droplet generation presented in **Fig. S4, S5**). The device is robust to translation across different droplet size regimes, and easy to adopt in different workflows. Alternative commercial devices or different device geometries are also compatible with sdDE-FACS.

### 3. Double Emulsion Device Wettability

To generate differential wettability between the first and second flow focuser of the ‘one step’ dual-flow focuser droplet generator device, we employ selective air or oxygen plasma treatment (4.5 min, 150W O_2_ plasma) on the inlets associated with the outer aqueous flow focuser (outer aqueous sheath inlet and outlet) using the tape method described previously4. Alternative surface treatments can be employed, if desired. Resultant double emulsions need only be relatively uniform (CV: <20%) and lacking significant free oil (by employing sdDE-FACS surfactant suggestions and adjusting flow rates).

### 4. Double Emulsion Surfactants

A table of surfactants used for sdDE-FACS is presented in **Table 1**. Any base buffer (***e.g.*** PBS, water, media, etc.) can be used for these surfactant formulations with no adverse effects as long as serum and other surfactants are removed or minimized. Optionally, BSA (0.5-2%) can be substituted for inner core Tween-20 (0.1-1%) with no adverse effects; this substitution may be desirable for cellular studies where viability is important.

### 5. Droplet Dilution Extended Note

Population ‘purity’ during sorting (***i.e.*** the percentage of events in the DE FSC ***vs.*** SSC gate) depends strongly on the absolute number droplets loaded into FACS diluent buffer (total volume: 600 uL), as highlighted in SH800 (Sony) data with highly permissive threshold and gain settings (**Fig. S2)**. Events in the lower left corner of this FSC ***vs.*** SSC plot indicate dust and small surfactant micelles that comprise ^~^38% of the parental population (with DEs comprising the remaining 61.6%, 30,000 total events in parental population on both Aria II and SH800). These results are typical for runs in which 50 – 100 μL are loaded from the droplet pellet (^~^0.2 – 5M droplets, dependent on size). Loading smaller numbers of DEs reduces overall throughput and may increase the need to manually resuspend DEs during a FACS run. Conversely, loading larger numbers of DEs can increase breakage through elastic collisions between DEs (particularly on the SH800) and therefore requires careful monitoring of droplet singlet rates over time. Events/second rates > 2,000 generally signify breakage and that the sort pressure should be decreased or the run should be aborted. In any case, subsequent gating allows stringent isolation of DEs from contaminants, rendering sdDE-FACS compatible with both scarce biological samples and abundant samples containing rare variants of interest.

### 6. Droplet Lag Time

On the SH800, after the start of a run there is typically a ‘lag’ time prior to the appearance of double emulsion droplets as shown in **Fig. S3**. On average, DEs are observed within the relevant FSC vs. SSC gate by 20s for the Aria II but can take up 1 – 4 min for the SH800, with a strong dependence on initial loading droplet density, droplet size and sort pressure. In order to boost initial events in the SH800, we begin all flow cytometry double emulsion runs at 9 psi until events begin to appear in the DE gate; after their appearance, droplet events tend to rise exponentially (as shown in **Fig. S3**). Subsequently, we modulate the sort pressure between 4 – 9 psi, corresponding to droplet density, to cap the event rate below 200 events/s. The SH800 uses a microfluidic chip with a laminar flow regime rather than the quartz cuvette with a hydrodynamic flow focusing regime employed by the Aria II. Observed droplet event delay and subsequent rapid boosting may be due to droplet packaging within the microfluidic sort chip or sample line. Double emulsions are highly deformable; similar effects have been observed and utilized for hydrogel packing regimes in microfluidics to achieve highly-regular flow metering after packing.

### 7. BD Aria II: Flow Cell Effects

The Aria II can be operated with a rectangular or square flow cell. Square flow cells observe “plug flow” behaviour and have excellent performance for large cell samples. sdDE-FACS performs well with either flow cell; a mixed population enrichment experiment is highlighted in **Fig. S8** using the square flow cell. All other experiments reported in this work were conducted with a rectangular flow cell.

### 8. SH800: Size Differentiation Effects

DEs can be easily discriminated by overall droplet size using the sdDE-FACS technique as shown in **Figs. S4, S5,** and **S6**. Size discrimination between 30 μm and 40 μm droplet populations is most apparent on FSC-A ***vs.*** SSC-A plots. Within populations, large droplet sizes (40 μm) can be more readily differentiated by inner volume and oil shell thickness (**Fig. S6**), especially using the SSC-A parameter which is able to parse 3 distinct scatter distributions corresponding to the 3 oil shell sizes shown in **Fig. S5**. This discrimination is pulse-width dominant (FSC-W and SSC-W effects are most significant in FSC-A and SSC-A discrimination).

---

**Figure S1:**
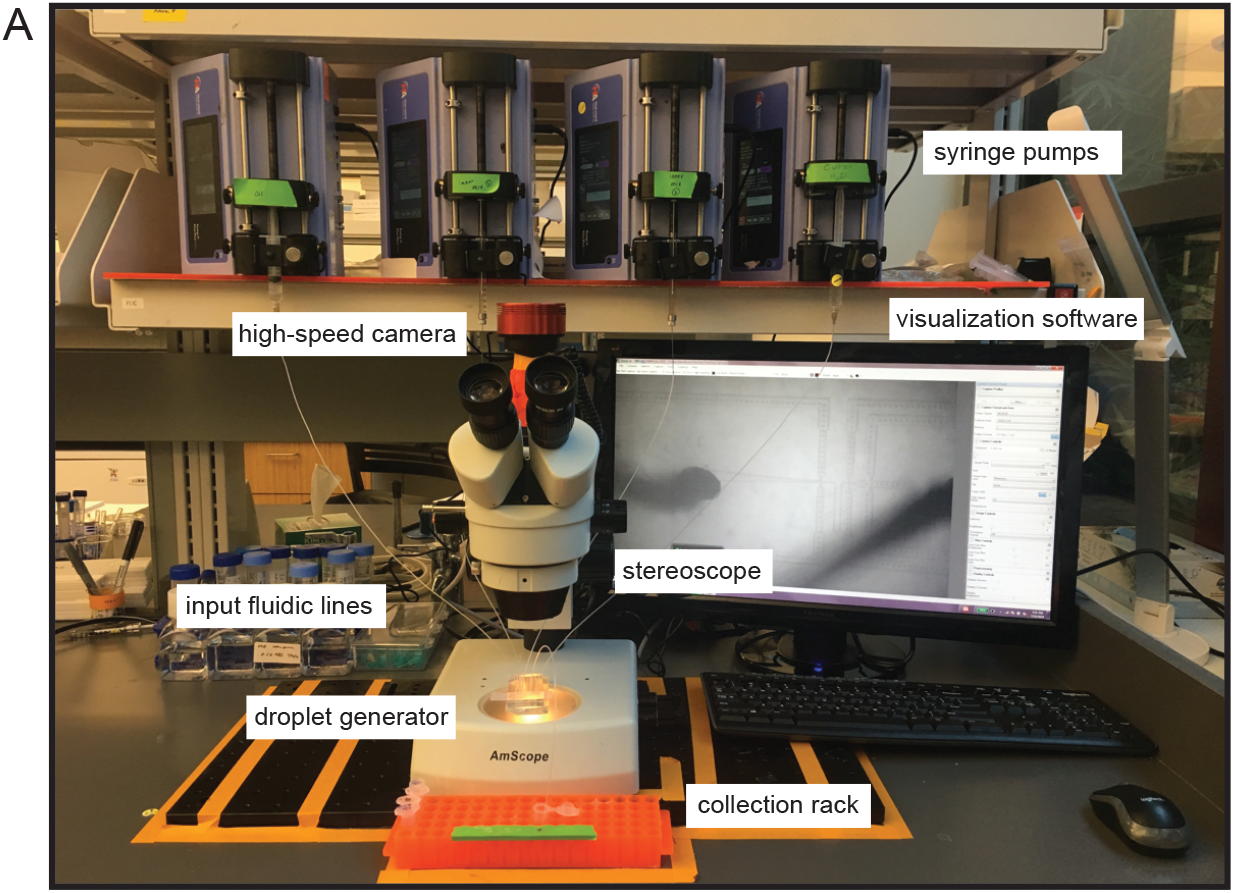
Double emulsion generation setup for sdDE-FACS workflow. **(A)** Double emulsion generation setup (^~^$10,000) comprised of syringe pumps, a microfluidic droplet generator device, a stereoscope and high-speed camera for droplet visualization, and a collection rack for holding generated droplets. Syringes loaded with reagents are connected to device inlets via polyethylene (PE) tubing, and device outlet is connected to a collection tube via a short segment of PE tubing to collect droplets.

**Figure S2:**
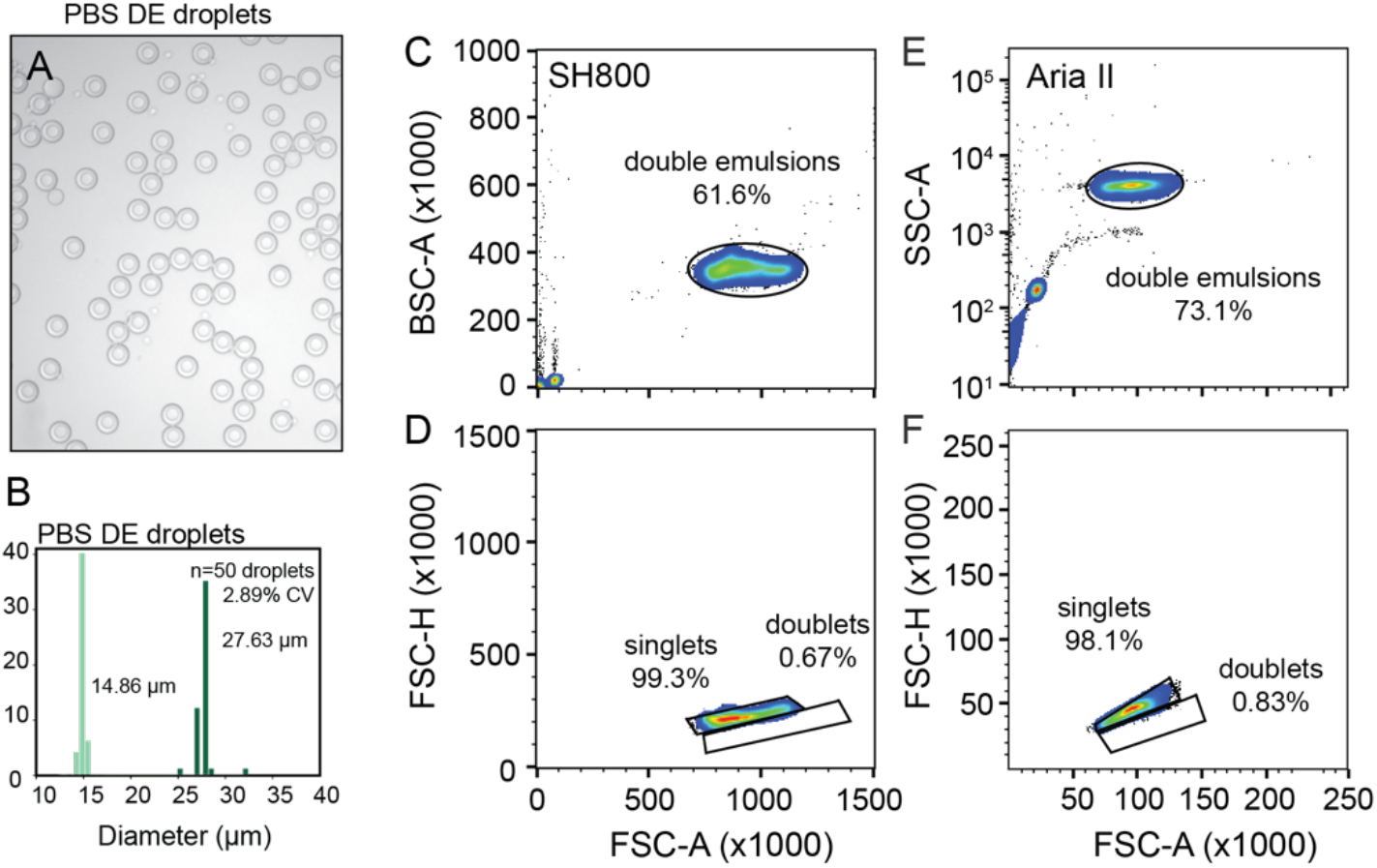
Representative sdDE-FACS analysis results for 30 μm droplets. **(A)** Brightfield image of DE droplets loaded with PBS buffer. **(B)** Population size histogram (light green = inner diameter, dark green = outer diameter, mean diameters indicated). **(C, D)** FACS morphology and singlet discrimination gates for the SH800. **(E, F)** FACS morphology and singlet discrimination gates for the Aria II. 30,000 events are shown for each parental gate; events were randomly downsampled.

**Figure S3:**
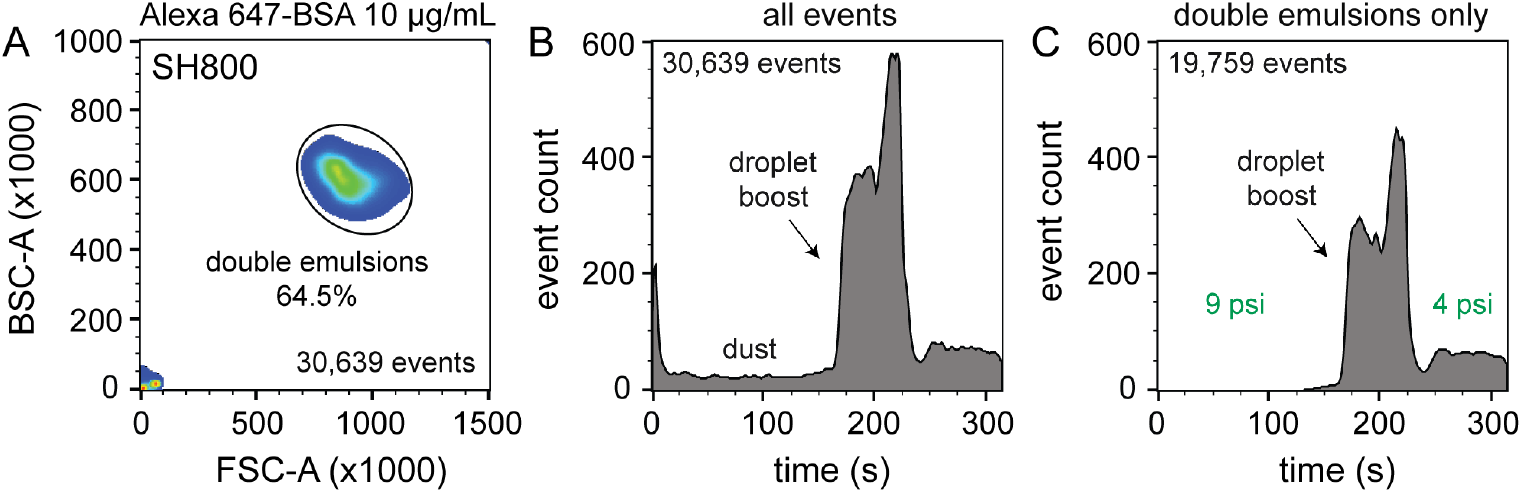
Typical delay times before the appearance of DE droplets on the SH800 (Sony). **(A)** Representative DE population as shown on the FSC ***vs.*** BSC morphology gate (SH800). **(B)** Total events per time shown for the full parental population of (A). **(C)** Events appearing in the double emulsion gate shown in (A). Note that dust events (shown in the lower left corner of (A)) dominate for the first ^~^160s of the FACs run. After ^~^160s, DE droplets begin to appear and event rates rise rapidly. Pressure was run at 9 psi (instrument limit = 10 psi per 130 μm nozzle) until the droplet event boost was observed. When a significant droplet population was visualized for gating, sheath pressure was decreased to 4 psi to attain a sort rate below 200 events/s for optimal droplet integrity post-sort.

**Figure S4:**
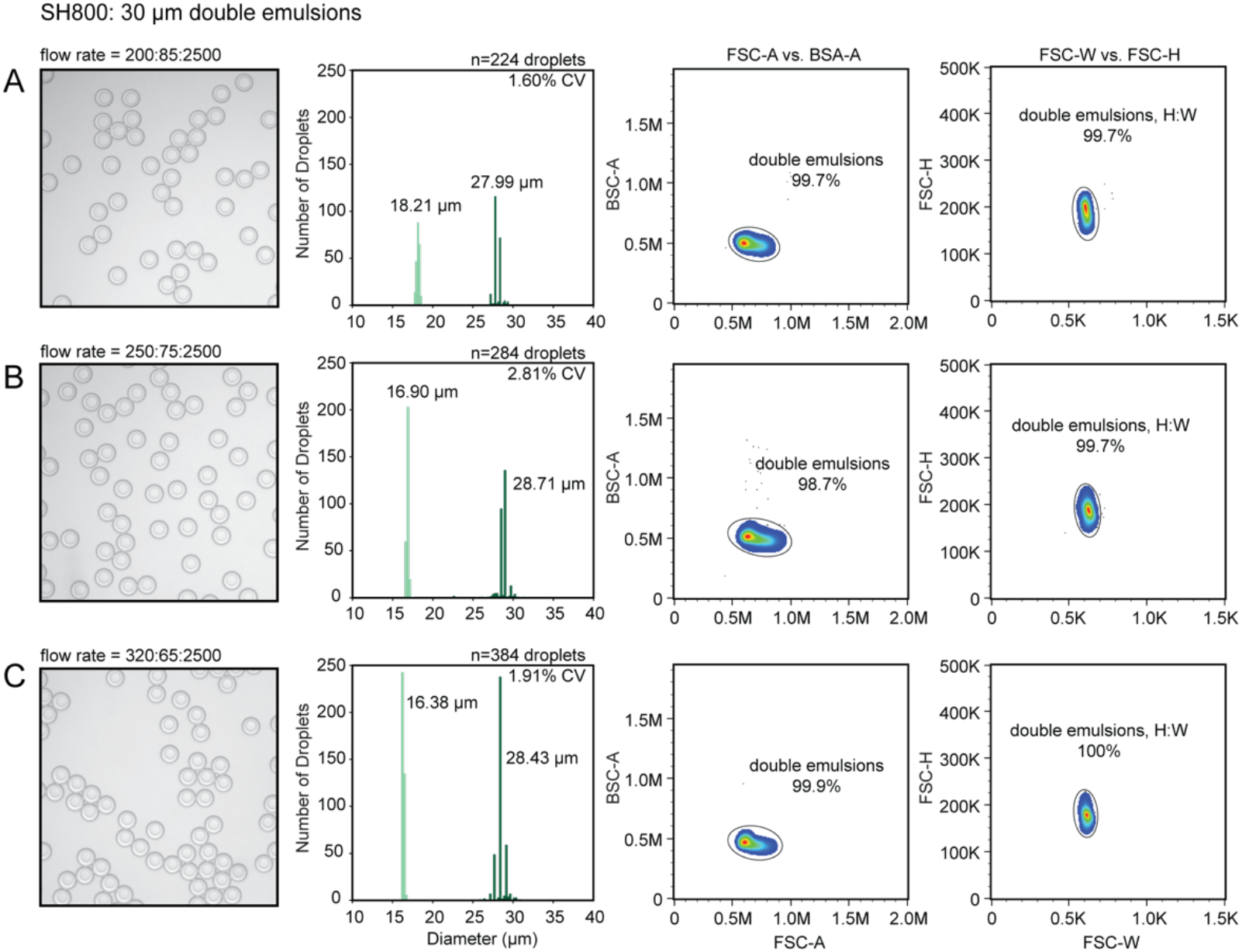
Size discrimination using sdDE-FACS on the SH800 (Sony) sorter, **Part I**. Comparative FACS analysis of 30 μm DE droplets with 3 different oil shell thickness resulting from different droplet flow rates (oil: inner core: outer aqueous sheath) of **(A)** 200:85:2500, **(B)** 250:75:2500, and **(C)** 320:65:2500. Representative brightfield images (left) and size histograms (light green = inner diameter, dark green = outer diameter, middle left) show population monodisperity and size differences between conditions. Ratio of inner core volume to total volume of the droplet are as follows for each condition: **(A)** 0.45, **(B)** 0.30, and **(C)** 0.24. FSC-A vs. BSC-A (middle right) and FSC-H vs. FSC-W (right) FACS plots for each condition (downsampled randomly to 1500 events per population) demonstrate that oil shell size only has minor effects on differential FSC or BSC in the 30 μm droplet populations.

**Figure S5:**
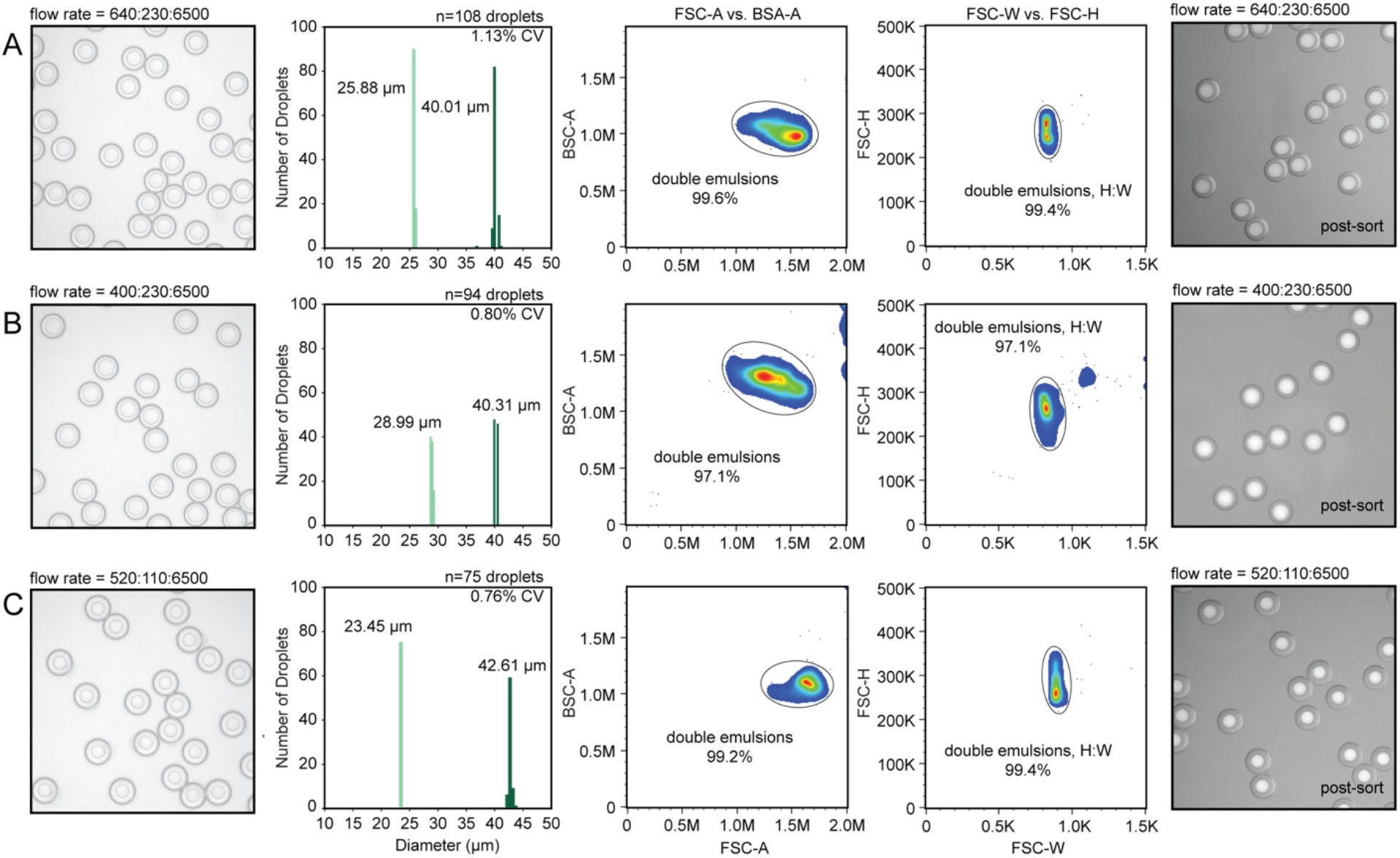
Size discrimination using sdDE-FACS on the SH800 (Sony) sorter, **Part II**. Comparative FACS analysis of 40 μm DE droplets with 3 different oil shell thickness resulting from different droplet flow rates (oil: inner core: outer aqueous sheath) of **(A)** 640:230:6500, **(B)** 400:230:6500, and **(C)** 520:110:6500.. Representative brightfield images (left) and size histograms (light green = inner diameter, dark green = outer diameter, middle left) show population monodisperity and size differences between conditions. Ratio of inner core volume to total volume of the droplet are as follows for each condition: **(A)** 0.37, **(B)** 0.59, and **(C)** 0.20. FSC-A vs. BSC-A (middle right) and FSC-H vs. FSC-W (right) FACS plots for each condition (downsampled randomly to 1500 events per population) demonstrate that oil shell size has large effects on differential FSC or BSC in the 40 μm droplet populations.

**Figure S6:**
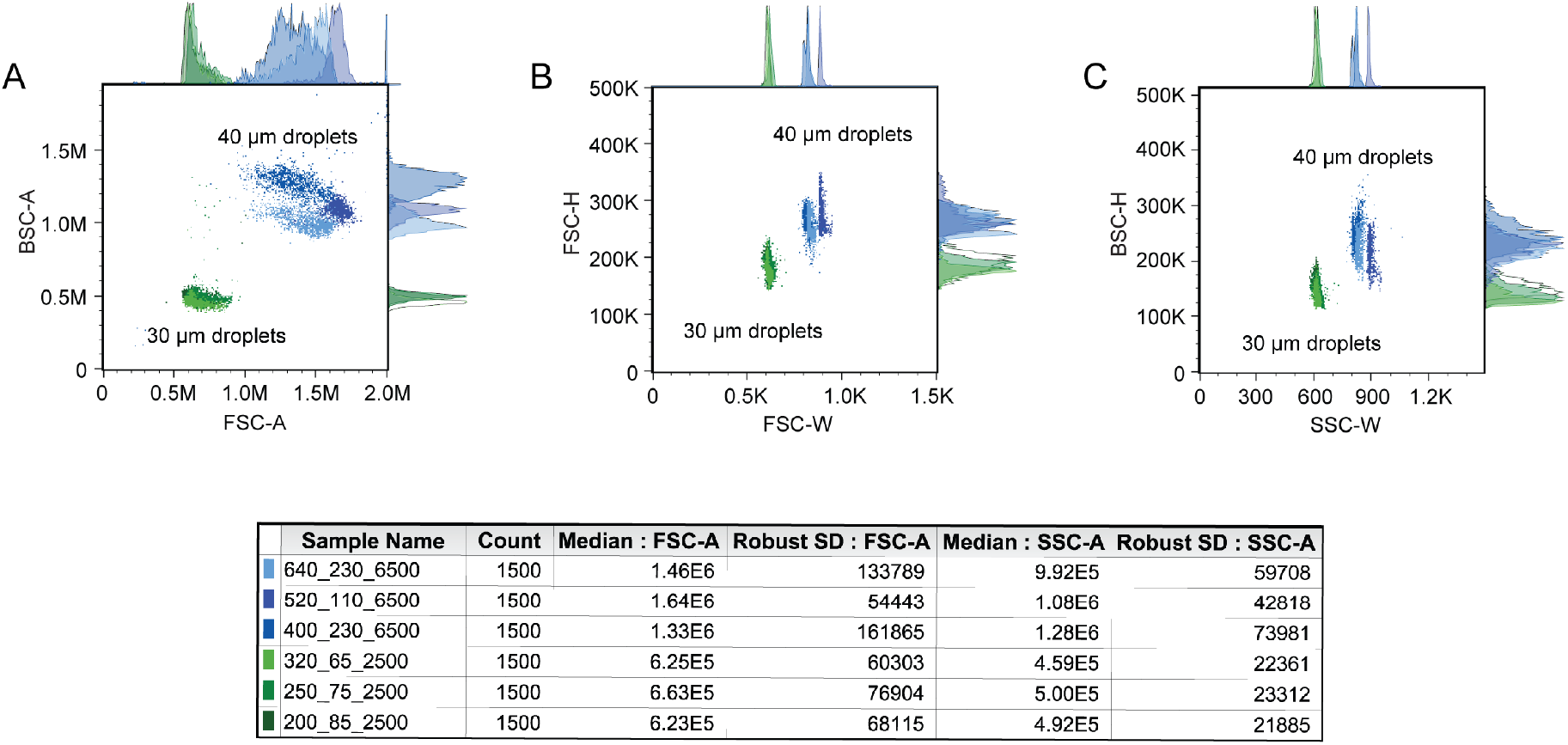
Comparative analysis of sdDE-FACS data 30 μm (**Fig. S4**) vs. 40 μm (**Fig. S5**) droplet populations. **(A)** FSC-A vs. BSC-A, **(B)** FSC-H vs. FSC-W, and **(C)** BSC-H vs. BSC-W distributions by condition reveal clear separation between 30 and 40 μm droplets and high inter-size discrimination on BSC-A for large droplets (40 μm population), with dominant effects from pulse-width discrimination. See further discussion in ***Extended Notes***.

**Figure S7:**
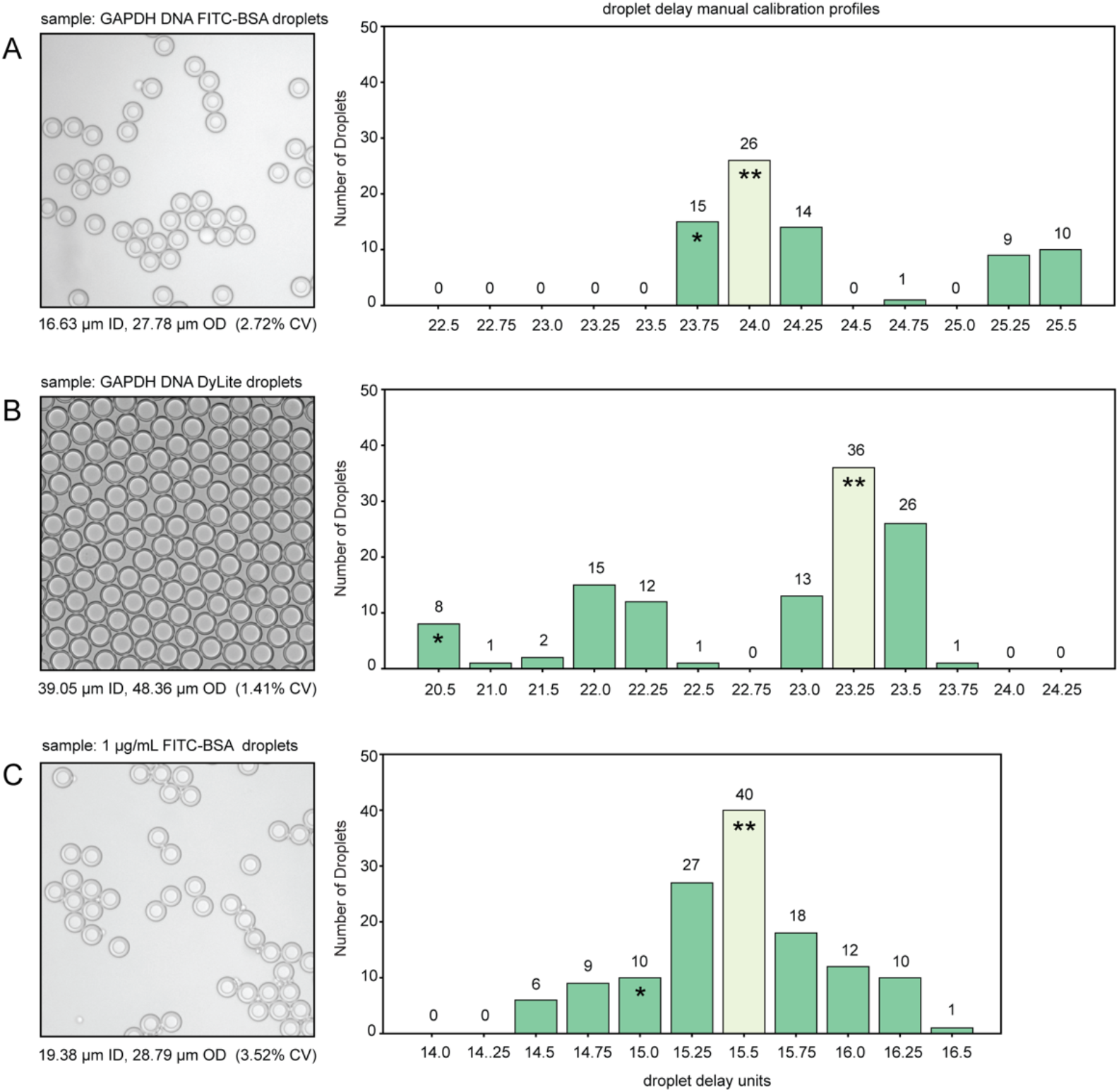
Representative droplet delay profiles from empirical droplet delay determination for 30-50 μm DEs on the Aria II. Manual delay profiles are shown for: **(A)** GAPDH-DNA loaded double emulsions with FITC-BSA dye (27.8 μm), **(B)** small-shell GAPDH-DNA loaded double emulsions with DyLite Antibody (Cy5) dye (48.4 μm), and **(C)** FITC-BSA reference droplets (28.9 μm). Each histogram was calculated for a manual droplet delay sweep outlined in ***Extended Methods.*** Each bar of the histogram corresponds to number of observed droplets in a well designated to receive 50 DE droplets at the indicated droplet delay. Accudrop-calibrated droplet delays and empirically determined drop delays are denoted by (*) and (**), respectively.

**Figure S8:**
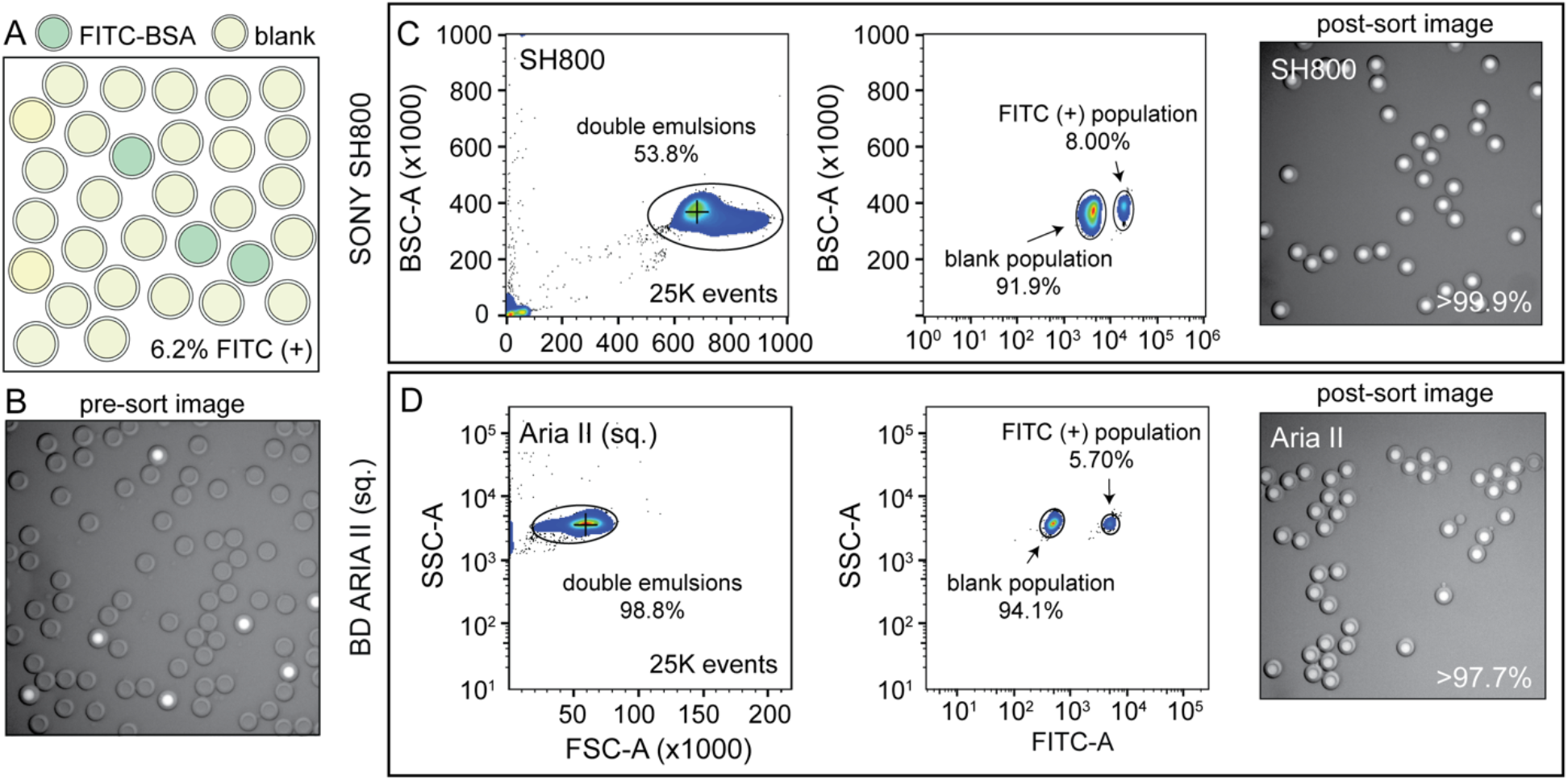
Replicate target enrichment experiment from a mixed parental population similar to **Fig. 4** conducted a square flow cell on the Aria II with manually-adjusted droplet delay. FITC-positive droplets were enriched with high specificity from a mixed population containing of 6.2% FITC-BSA positive DEs and blank DEs **(A, B).** High target specificity and sensitivity were observed on an Aria II instrument containing a square flow cell, comparable to rectangular flow cell results (**Fig. 4**). See further discussion in ***Extended Notes***.

**Figure S9:**
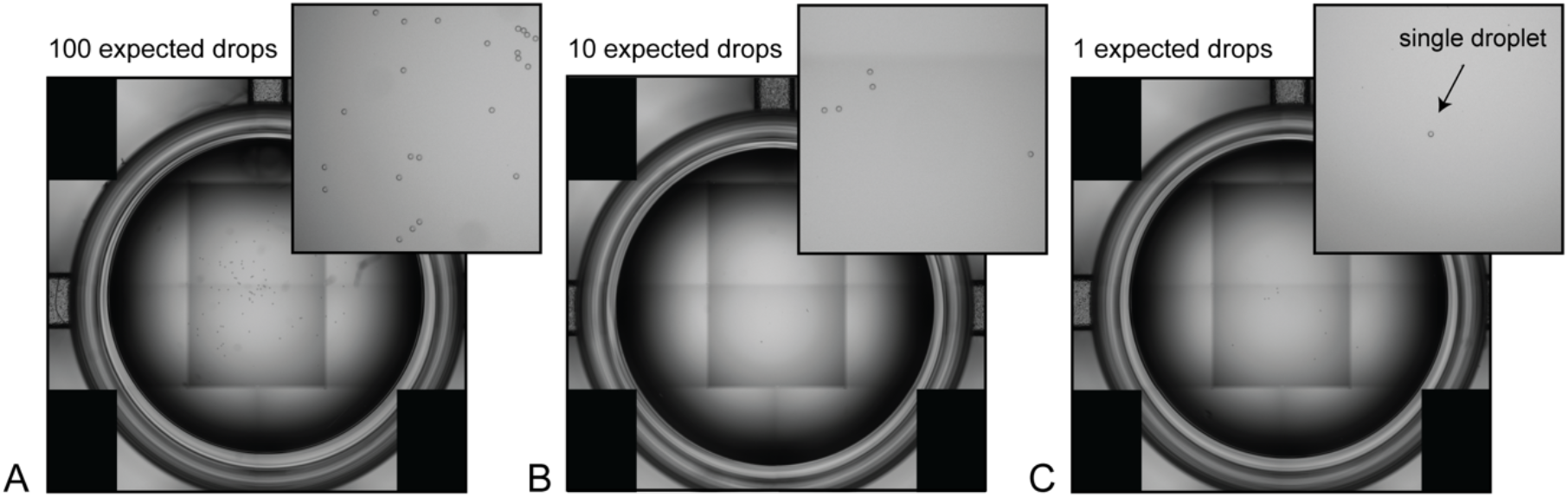
Example EVOS (Life Technologies) microscopy images used to calculate DE-FACS recovery rates using the Aria II sorter for **(A)** 100-, **(B)** 10-, and **(C)** 1-droplet set points. Each panel displays a full-well image (left) and magnified 600 × 600 pixel region of interest (top right).

**Figure S10:**
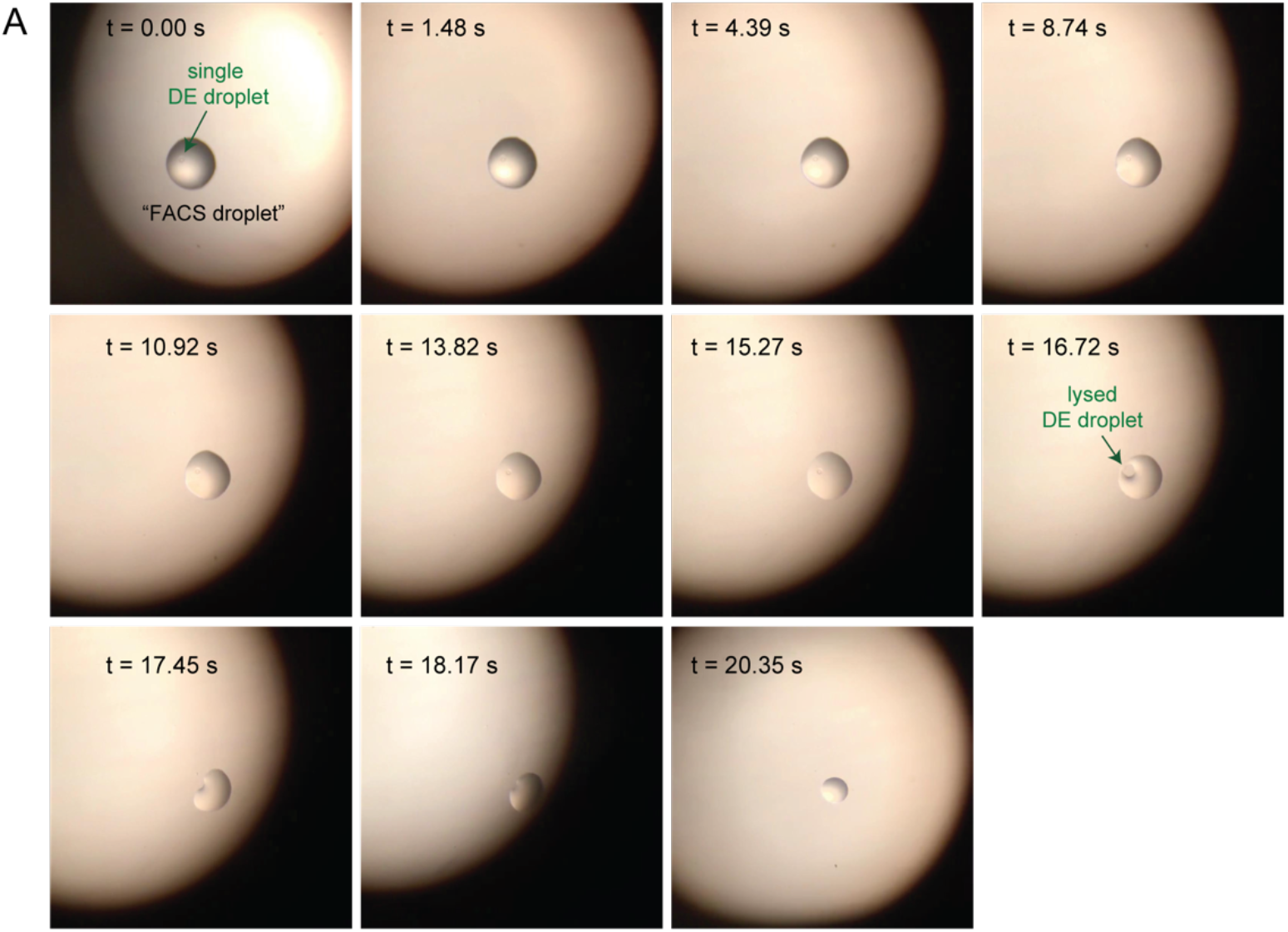
Image sequence of on-plate droplet lysis within a single ‘FACS droplet’ deposited in a dry well of a 96-well plate. Dry droplet lysis was employed for nucleic acid recovery as presented in **Figs. 6, S11**.

**Figure S11:**
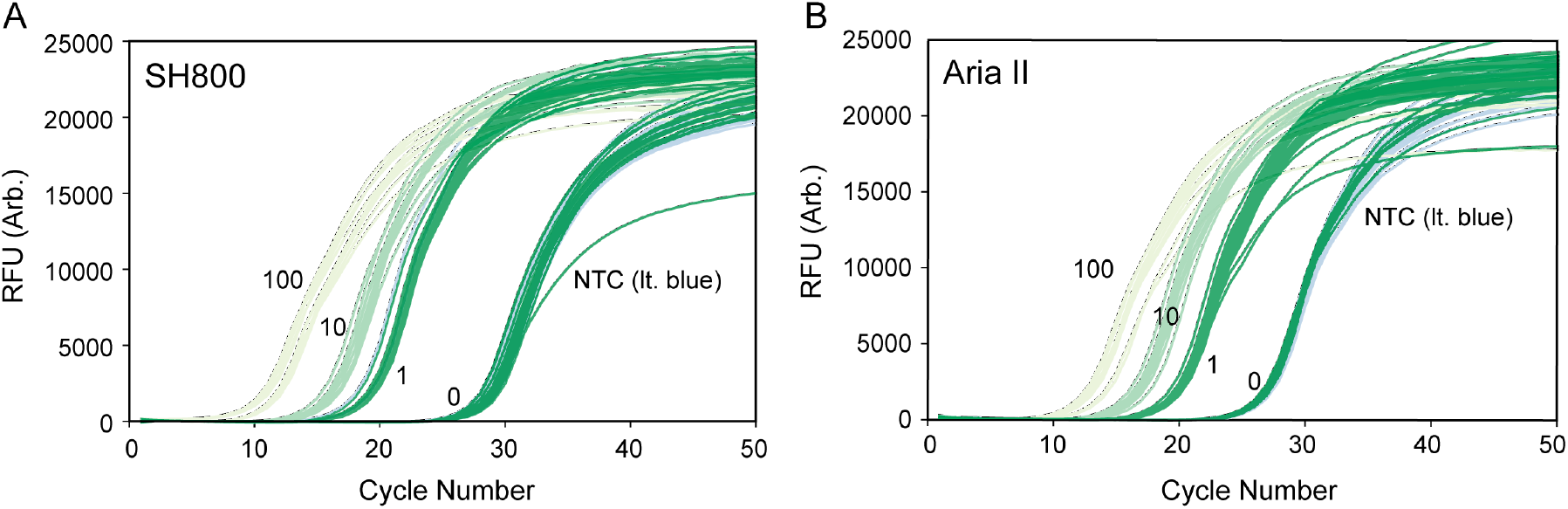
Raw qPCR traces using sdDE-FACS with two flow cytometer instruments: **(A)** SH800 (Sony), and **(B)** Aria II. qPCR traces are for 96-well full-plate nucleic acid recovery (standards omitted).

**Figure S12:**
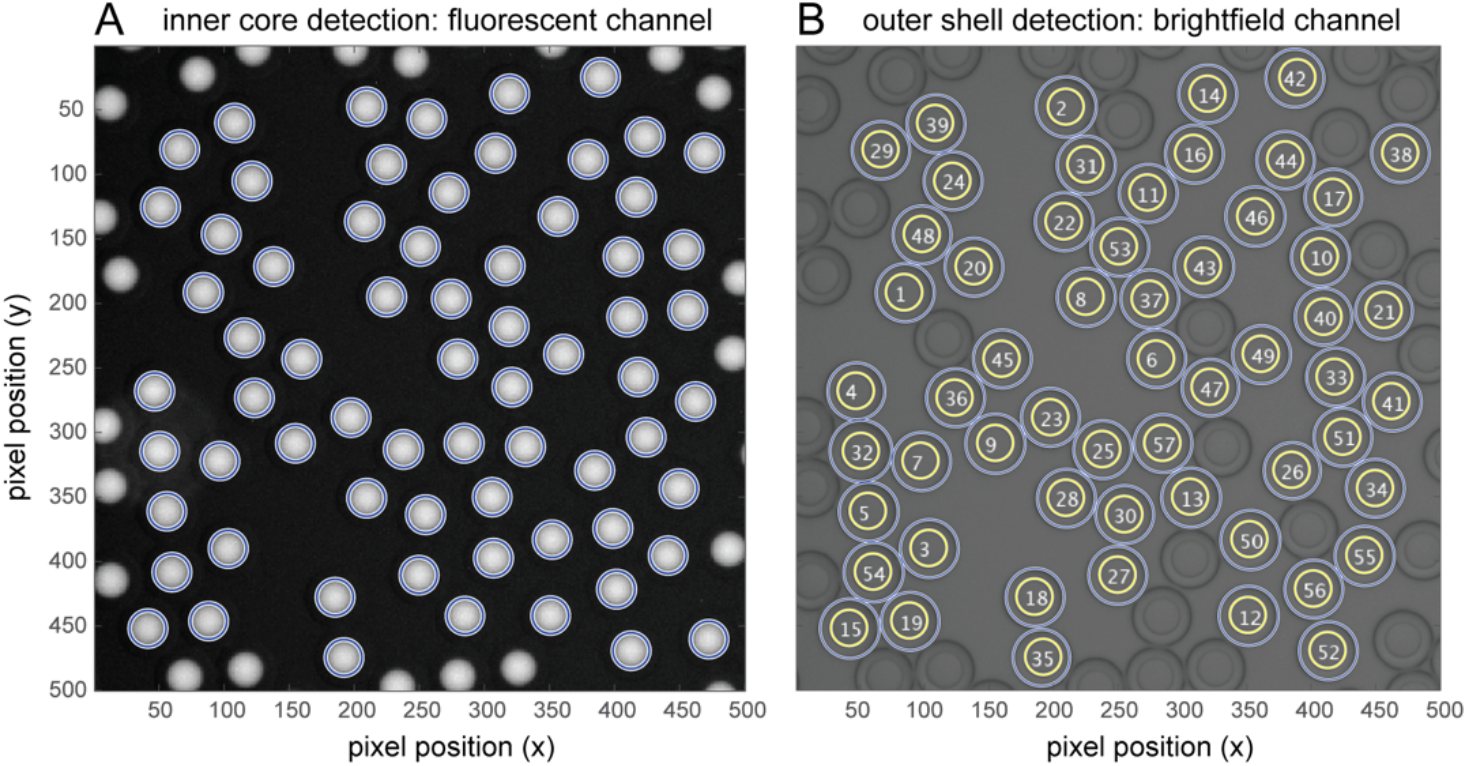
Automated DE droplet detection and inner and outer droplet size measurement. **(A)** A modified Hough transform algorithm allows automated droplet identification and measurement of inner core diameters from a reference dye fluorescent channel image, as shown here for DE droplets containing FITC-BSA. **(B)** Outer shell boundaries are identified by performing a line scan centered on the fluorescence centroid position and searching for intensity minima in the greyscale image. Droplets are indexed by condition, image number, and droplet count, as shown; overlapping or adjacent droplets are omitted from subsequent size analysis.

**Table S-T1:** Components and associated suppliers for building and operating the droplet generation setup shown in **Figure S1**. Note: the syringe pumps listed (Pico Pump Elite, Harvard Apparatus) are utilized specifically for high-precision, low-volume applications (10 – 500 μL/hr); alternative suppliers can be used (New Era Pumps, $1,500/pump) to reduce costs if desired. Costs current as of October 2019.

| Setup Component | Supplier | Cost |
| --- | --- | --- |
| Syringe pumps | Pico Pump Elite, Harvard Apparatus | \$3,083.00 x 3 |
| High-frame rate camera | ASI 174MM, ZWO ASI | \$599.00 |
| Visualization software | SharpCap, ZWO ASI | \$0.00 |
| Computer | Generic laptop, Lenovo | \$650.00 |
| PDMS Device | RTV 615, Momentive (*per chip*) | \$3.00 |
| Microfluidic tubing | PE/2, Sci. Commodities (*per spool*) | \$83.00 |
| Syringes, disposable | 1 mL, 5 mL Plasti-pak, BD (*per run*) | \$6.00 |
| Total | | \$10,590.00 |

**Table S-T2:**
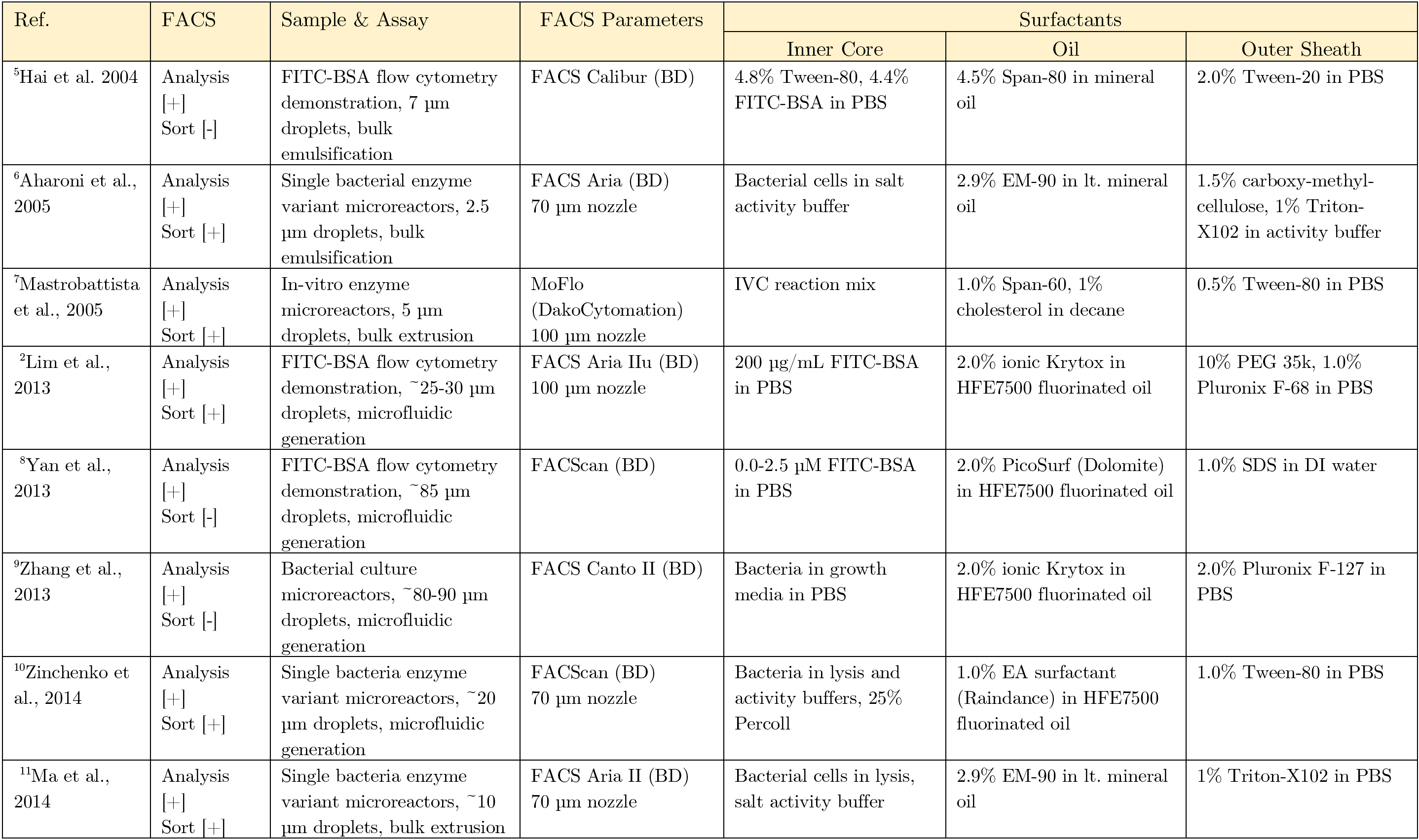
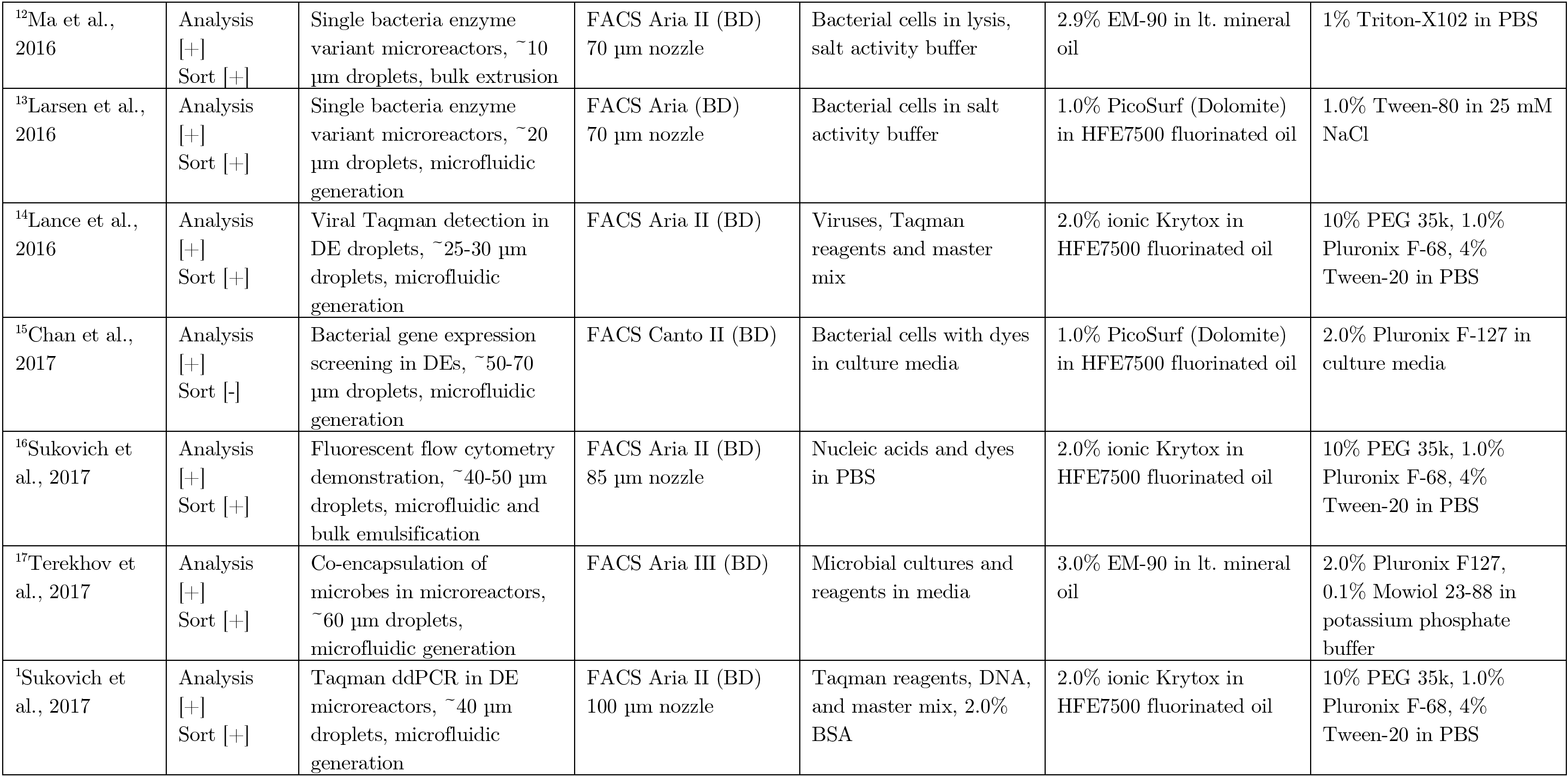
Reported surfactant mixtures and FACS cytometer settings for prior literature in double emulsion flow cytometric analyses. Nozzle sizes are listed when reported or obtained from authors. Surfactant formulations compare directly to those presented in **Table 1** for sdDE-FACS.

**Table S-T3:** Extended plate statistics. Manual droplet delay calibration is indicated by (*). qPCR readouts contain 100, 10, 1 and NTC wells as indicated in **Figs. 6, S11**; however, binary sort statistics corresponding to droplet presence of absence (as determined by C_q_ cluster and expected [DNA]) are only accessible for single droplet deposition wells (n=36 wells/plate) and are reported here as fraction of total occupied single-droplet set point wells.

| Assay | Sorter | Sample | Set Point (Expected No. of Droplets) |  |  |  |
| --- | --- | --- | --- | --- | --- | --- |
|  |  |  | 100 droplets | 10 droplets | 1 droplet | 0 droplet |
| Optical | Aria II* | GAPDH DyLite | 71.2 (5.6)<br>N=11 wells | 6.1 (2.2)<br>N=36 wells | 0.50 (0.51)<br>N=24 wells | 0 (0)<br>N=25 wells |
| Optical | SH800 | GAPDH DyLite | 69.9 (4.5)<br>N=12 wells | 7.1 (1.5)<br>N=36 wells | 0.71 (0.50)<br>N=24 wells | 0 (0)<br>N=24 wells |
| Optical | Aria II* | Plate 1<br>FITC-BSA | 81.9 (4.0)<br>N=12 wells | 6.9 (3.5)<br>N=12 wells | 0.83 (0.38)<br>N=48 wells | 1 (0)<br>N=24 wells |
| Optical | Aria II* | Plate 2<br>FITC-BSA | 83.6 (2.7)<br>N=12 wells | 8.4 (1.2)<br>N=12 wells | 0.83 (0.38)<br>N=48 wells | 0 (0)<br>N=24 wells |
| Optical | SH800 | Plate 1<br>FITC-BSA | 55.5 (3.4)<br>N=12 wells | 5.3 (1.8)<br>N=12 wells | 0.75 (0.48)<br>N=48 wells | 0 (0)<br>N=24 wells |
| Optical | Aria II* | Plate 1<br>GAPDH DyLite | 64.4 (7.9)<br>N=12 wells | 5.9 (1.7)<br>N=12 wells | 0.77 (0.47)<br>N=48 wells | 0 (0)<br>N=24 wells |
| Optical | Aria II* | Plate 2<br>GAPDH DyLite | 64.8 (7.2)<br>N=12 wells | 6.3 (1.7)<br>N=12 wells | 0.56 (0.61)<br>N=48 wells | 0 (0)<br>N=24 wells |
| Optical | SH800 | Plate 1<br>GAPDH DyLite | 46.2 (8.2)<br>N=12 wells | 4.9 (1.4)<br>N=12 wells | 0.56 (0.50)<br>N=48 wells | 4 (0)<br>N=24 wells |
| Optical | Aria II* | *Singles-Only*<br>GAPDH DyLite | 0 (0)<br>N=0 wells | 0 (0)<br>N=0 wells | 0.80 (0.40)<br>N= 96 wells | 0 (0)<br>N=0 wells |
| qPCR | SH800 | Plate 1<br>DNA-FITC-BSA | N/A (--)<br>N=0 wells | N/A (--)<br>N=0 wells | 0.53 (--)<br>N=19/36 wells | N/A (--)<br>N=0 wells |
| qPCR | SH800 | Plate 2<br>DNA-FITC-BSA | N/A (--)<br>N=0 wells | N/A (--)<br>N=0 wells | 0.69 (--)<br>N=25/36 wells | N/A (--)<br>N=0 wells |
| qPCR | Aria II* | Plate 1<br>DNA-FITC-BSA | N/A (--)<br>N=0 wells | N/A (--)<br>N=0 wells | 0.61 (--)<br>N=22/36 wells | N/A (--)<br>N=0 wells |
| qPCR | Aria II* | Plate 1<br>DNA-FITC-BSA | N/A (--)<br>N=0 wells | N/A (--)<br>N=0 wells | 0.66 (--)<br>N=23/35 wells | N/A (--)<br>N=0 wells |

**Table S-T4:**
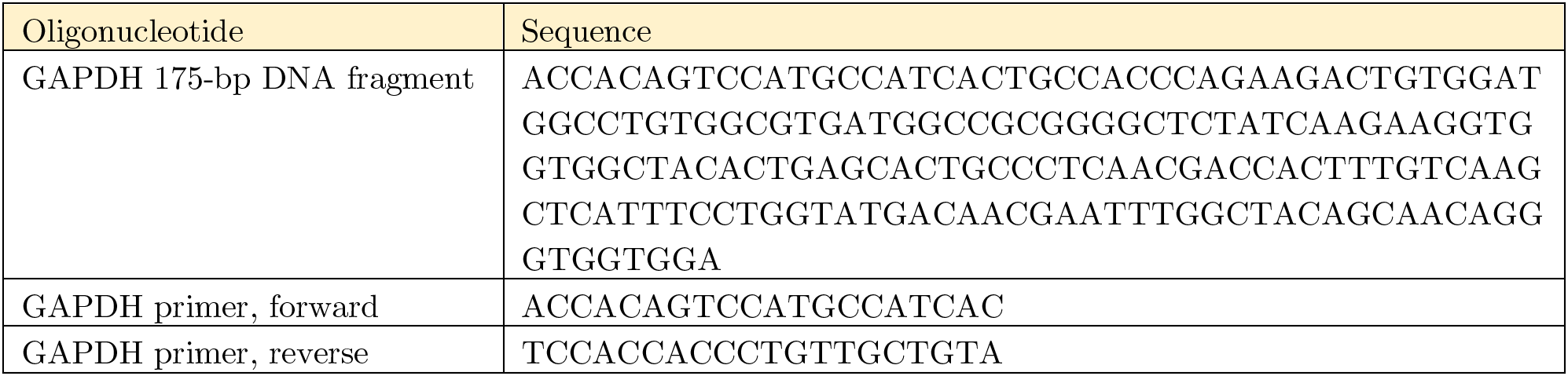
Oligonucleotides used for DE nucleic acid recovery qPCR experiments described in **Fig. 6** and **Fig. S11**.

**Table S-T5:** Reaction components for qPCR experiments described in **Fig. 6** and **Fig. S11.**

| Reaction Component | Volume *per well* | Final Concentration |
| --- | --- | --- |
| 2X iTaq SYBR qPCR Mix (Biorad) | 5 uL, 1X | 1X |
| 10 µMGAPDH primer, forward | 0.25 uL | 250 µM |
| 10 µMGAPDH primer, reverse | 0.25 uL | 250 µM |
| Nuclease-free Water | 4.5 µL | - |
| Total | 10 µL | - |

